# Protein surface chemistry encodes an adaptive tolerance to desiccation

**DOI:** 10.1101/2024.07.28.604841

**Authors:** Paulette Sofía Romero-Pérez, Haley M. Moran, Azeem Horani, Alexander Truong, Edgar Manriquez-Sandoval, John F. Ramirez, Alec Martinez, Edith Gollub, Kara Hunter, Jeffrey M. Lotthammer, Ryan J. Emenecker, Hui Liu, Janet H. Iwasa, Thomas C. Boothby, Alex S. Holehouse, Stephen D. Fried, Shahar Sukenik

## Abstract

Cellular desiccation - the loss of nearly all water from the cell - is a recurring stress in an increasing number of ecosystems that can drive protein unfolding and aggregation. For cells to survive, at least some of the proteome must resume function upon rehydration. Which proteins tolerate desiccation, and the molecular determinants that underlie this tolerance, are largely unknown. Here, we apply quantitative and structural proteomic mass spectrometry to show that certain proteins possess an innate capacity to tolerate rehydration following extreme water loss. Structural analysis points to protein surface chemistry as a key determinant for desiccation tolerance, which we test by showing that rational surface mutants can convert a desiccation sensitive protein into a tolerant one. Desiccation tolerance also has strong overlap with cellular function, with highly tolerant proteins responsible for production of small molecule building blocks, and intolerant proteins involved in energy-consuming processes such as ribosome biogenesis. As a result, the rehydrated proteome is preferentially enriched with metabolite and small molecule producers and depleted of some of the cell’s heaviest consumers. We propose this functional bias enables cells to kickstart their metabolism and promote cell survival following desiccation and rehydration.

**Teaser:** Proteins can resist extreme dryness by tuning the amino acids on their surfaces.

## Introduction

In a desiccating cell, proteins, ions, and small molecules experience a dramatic increase in concentration. Many proteins exist at the edge of their solubility limits, even under well-hydrated conditions (*1–3*). As this limit is reached, proteins become super-saturated leading to their aggregation and inactivation, often irreversibly. For an organism to survive desiccation, at least some of its proteome must continue to function upon rehydration (*4*). This can be facilitated by “priming,” the process by which cells take up or synthesize high levels of protective molecules during the onset of water loss (*5*, *6*). These include small polyols such as trehalose, sorbitol, and sucrose (*7*, *8*), as well as larger proteins such as heat shock proteins (e.g., Hsp20s (*9*)) and specialized disordered proteins (e.g., CAHS or LEA proteins (*10*, *11*)). However, some organisms can survive desiccation even without priming, and the intrinsic ability of proteomes to tolerate desiccation is poorly characterized (*12*, *13*).

Our understanding of the molecular consequences of desiccation is largely based on reconstitution assays, examining the impact on specific proteins under well-controlled conditions (*14*). Biochemical assays examining enzymatic activity before and after desiccation have provided evidence that many proteins lose their function via aggregation or misfolding (*15–18*). However, these findings are anecdotal and focus on the loss of activity for a small number of proteins, often in purified form and under dilute buffer conditions. While these experiments benefit from being well-controlled, they do not necessarily reflect the heterogeneous cellular milieu (*19*). To uncover the molecular mechanisms behind successful protein rehydration, we considered an assay that could provide a holistic examination of the entire proteome.

Here, we used mass spectrometry to study how a proteome withstands desiccation and rehydration. Using undiluted lysates obtained by cryogenic pulverization of *Saccharomyces cerevisiae*, we quantified the resolubility of thousands of proteins. This was done by measuring the change in abundance of soluble lysate proteins following desiccation and subsequent rehydration (desiccation-rehydration; D-R). We find that, on average, proteins partition 1:3 between the supernatant (resoluble) and pellet (non-resoluble) fractions, with large deviations from this average for individual proteins. Additionally, using limited proteolysis mass spectrometry (LiP-MS) (*20*, *21*), we determined that resolubility and structural retention following D-R are highly correlated

To understand the chemical and structural determinants of protein desiccation tolerance, we correlated our results with structural predictions of the *S. cerevisiae* proteome (*22*, *23*). These analyses revealed that smaller, well-folded proteins with fewer disordered regions and fewer known protein interactors tend to have a high tolerance to D-R. Tolerance also correlated strongly with specific chemical properties of the protein surface: an enrichment of negative (but not positive) charged as well as small (Gly/Ala/Pro) residues.

Strikingly, the attributes that promote desiccation tolerance occur preferentially in proteins with specific functions: tolerant proteins are largely responsible for biosynthesis of small molecules and metabolites. Conversely, non-tolerant proteins are those that consume large portions of the cell’s resources, such as ribosome biogenesis (*24*). We propose that this enrichment of resource producers and depletion of consumers is an adaptive mechanism that kickstarts cellular metabolism upon rehydration and facilitates cell survival.

### The unprimed yeast proteome is poorly resoluble

To understand proteins’ intrinsic desiccation tolerance, we aimed to profile the proteome in the absence of protective molecules that are upregulated in response to stress and are known to confer global desiccation tolerance (*25*, *26*). During logarithmic growth phase, *S. cerevisiae* cells do not biosynthesize or uptake such molecules (*27*), significantly reducing their desiccation resistance compared to cells in the stationary phase (**Fig. 1A**) (*7*, *27*). We subjected *S. cerevisiae* cells, grown to mid-log phase, to cryogenic milling without any lysis buffer, which yielded an undiluted lysate containing cellular solutes and proteins near their physiological concentrations in the cell (*28*) (**Fig. 1B**). The lysate was subjected to a single cycle of desiccation-rehydration (D-R) (**Fig. 1C**). The total neat lysate sample (**T**) was vacuum dried in a desiccator until sample weight plateaued (**Fig. 1D**). Thermogravimetric measurements revealed that the desiccated lysate’s water content was 11.2%, in line with previous definitions for a desiccated state (*29*) (**Fig. 1D**).

**Figure 1.**
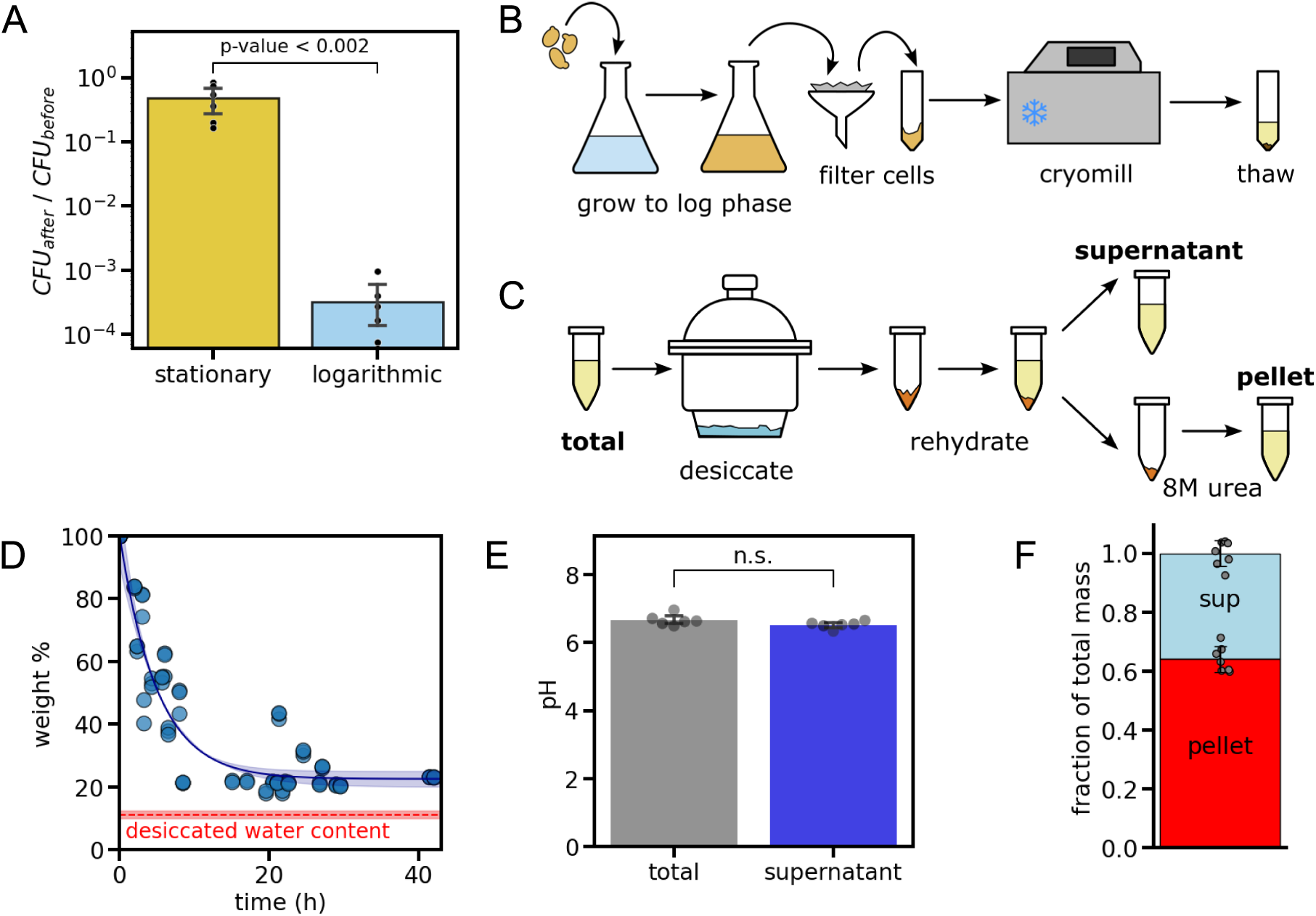
Neat lysates characterization before and after desiccation. (**A**) Surviving fraction of colony-forming units (CFUs) of stationary and log-phase yeast based on dilution plating from liquid cultures before and after desiccation-rehydration (D-R). Error bars are the standard deviation of 6 independent repeats. (**B**) Schematic of the lysate preparation procedure. (**C**) The desiccation-rehydration (D-R) process used to isolate proteins before (**total**, **T**) and after D-R into soluble (**supernatant**, **S**) or insoluble (**pellet**, **P**) fractions. (**D**) Lysate weight (in %) over the course of desiccation. Blue points are each an average taken from three independent treatments, with a line showing an exponential decay fit to the data. The dashed red line shows the total water content as determined by TGA after 40 hours. (**E**) pH of the soluble lysate before and after desiccation. Points were obtained from three independent samples, and error bars are standard deviations. (**F**) Protein mass distribution between the supernatant and pellet following desiccation and rehydration. Points were obtained from eight independent measurements, and error bars are standard deviations.

Following desiccation, dried lysates were rehydrated with pure water, and the supernatant (**S**) was separated from the pellet (**P**) via centrifugation. The supernatant showed no change in pH compared to the initial lysate (**Fig. 1E**). Protein quantification using a bicinchoninic acid (BCA) assay revealed that ∼35% of the protein mass remained soluble, while the remainder aggregated (**Fig. 1F**).

### Protein resolubility correlates with structural fidelity

Next, we used liquid chromatography tandem mass spectrometry (LC-MS/MS) to evaluate the capacity of each protein to resolubilize following D-R by quantifying relative protein abundances in the total, supernatant, and pellet fractions across three biological replicates. We reliably quantified 4,384 proteins in all three biological replicates, representing about 75% of the whole *S. cerevisiae* proteome (**Fig. 2A, Table S1**). Protein abundances in the total, supernatant, and pellet fractions and their rank orders were highly similar across the replicates (**Fig. S1**). Protein resolubilities were calculated by dividing protein abundance in the supernatant by the abundance in the total sample (**S**/**T**), which also showed a high degree of reproducibility (**Fig. S2**). An independent metric of the same measure, **S**/(**S**+**P**), strongly correlated with **S**/**T**, suggesting that the mass balance for most proteins in our experiment approaches 100% (**Fig. 2B**). From here on, we utilize the **S**/**T** ratio as a measure of a protein’s resolubility.

**Figure 2.**
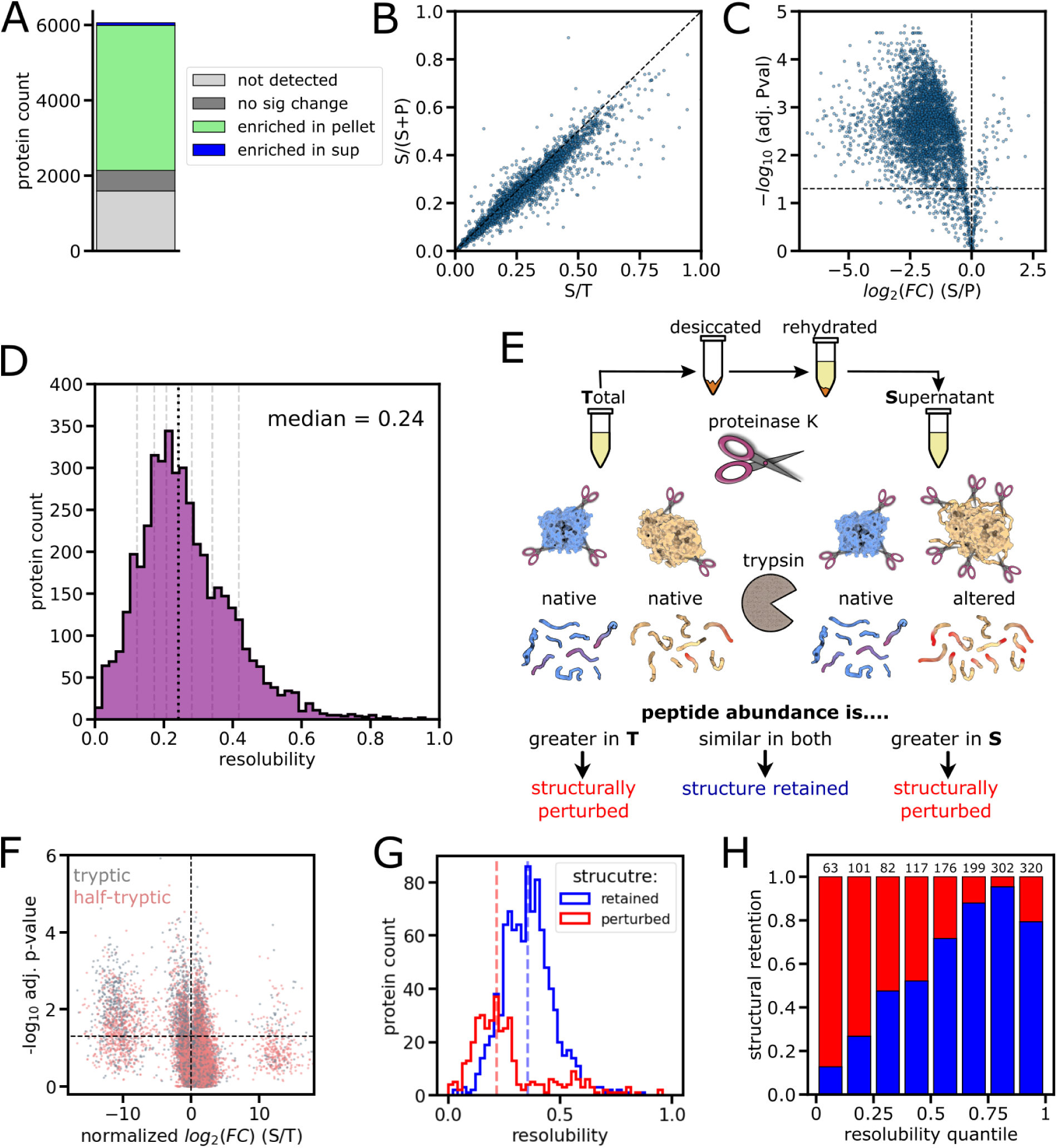
Proteome-wide resolubility and structure retention of the rehydrated proteome. **(A)** Proteome coverage of LC-MS profiling of lysates from *Saccharomyces cerevisiae* (UniProt UP000002311). (**B**) Correlation between S/(S+P) and S/T, two independent metrics for protein resolubility. Pearson’s r = 0.98. (**C**) Volcano plot of proteome partitioning between supernatant and pellet following D-R. Dashed horizontal line represents cutoff for statistical significance (FDR adjusted p-value < 0.05). (**D**) Histogram of proteome resolubility fractions. Grey dashed lines represent 1/8th quantiles (∼ 550 proteins in each quantile). Median resolubility is 0.24, shown as a black dashed line. (**E**) Schematic of experimental workflow for limited proteolysis mass spectrometry (LiP-MS). (**F**) Volcano plot showing the normalized relative abundance of full-tryptic and half-tryptic peptides before and after D-R. Horizontal line represents cutoff for significance (FDR adjusted p-value < 0.05). (**G**) Distribution of resolubility percentages for proteins whose structure was retained or perturbed. Dashed vertical lines are the medians. (**H**) Fraction of proteins that retained their structure for each resolubility quantile. Numbers on top indicate the number of proteins identified in each quantile by LiP-MS.

Overall, proteins partition primarily into the pellet following desiccation. Less than 2% of the proteins quantified (66/4384) partition preferentially into the supernatant significantly (adjusted p-value < 0.05), while over 88% (3855/4384) partition preferentially into the pellet significantly (**Fig. 2A**). In line with our BCA assay (**Fig. 1F**), the median resolubility amongst the 4,384 proteins assessed is 0.24, with a long tail towards higher values (**Fig. 2D**). However, this quantitative proteomics study does not reveal if the resolubilized proteins also retain their native structure after rehydration - which for many proteins is prerequisite for proper function.

To assess structural changes in rehydrated proteins following desiccation and rehydration, we employed limited proteolysis mass spectrometry (LiP-MS) on the resolubilized proteins (fraction **S**) and compared the proteolysis profile to extracts that were not subject to the D-R stress (fraction **T**). Briefly, **T** and **S** samples were incubated for one minute with the non-specific protease proteinase K. This protease preferentially cuts the protein at solvent-exposed regions, resulting in structural information that can be read out by sequencing the resulting peptide fragments with LC-MS/MS (*20*, *30*). The resulting peptides were further fragmented via trypsin digest and prepared for shotgun proteomics (**Fig. 2E**). The half-tryptic fragments – in which one cut arises from trypsin and another from proteinase K – indicate sites that were susceptible to proteolysis by proteinase K (i.e., solvent accessible regions); the full-tryptic fragments indicate regions in which proteinase K did not cut (i.e., buried regions). By identifying peptides with significant changes in abundance between **S** and **T** samples (normalized for overall protein abundance differences as measured by their resolubilities), we can assign sites within proteins that were structurally perturbed by the D-R cycle (**Fig. 2E**) (*20*).

The LiP-MS experiment (*31*) assessed the structural status for 1750 proteins, or 40% of the proteins with assigned resolubility (**Table S2**). In general, half-tryptic peptides were more abundant in **S** and full-tryptic peptides were more abundant in **T**. Specifically, 69% of the peptides that were >2-fold more abundant in **S** are half-tryptic, compared to 64% of peptides that had similar abundance in both (within 2-fold), and compared to 55% of peptides that were >2-fold more abundant in **T**. Hence, resolubilized proteins following D-R are generally more susceptible to proteolysis due to structural perturbation. (**Fig. 2F**). The number of identified half-tryptic and full-tryptic peptides were consistent across conditions and replicates, suggesting comparable proteolysis (**Fig. S3A-C**). Proteins were labeled structurally perturbed if they possessed two or more peptides with significantly different abundance (>2 fold-change between **S** and **T** following normalization for protein abundance, p < 0.05 by FDR-adjusted t-test with Welch’s correction) in the rehydrated supernatant. By this metric, 9.7% of peptides identified corresponded to sites that were significantly structurally perturbed, and 28.1% of identified proteins with more than one peptide mapped were structurally perturbed (**Fig. S3D-E**). The majority of peptides demonstrated consistency across biological replicates,, with a median coefficient of variation around 0.18 (**Fig S3F**).

We found that the resolubility distribution of proteins whose structure is perturbed is significantly different from that of proteins where the structure has been retained (p-value < 10^-58^ by KS 2-sided test) (**Fig. 2G, Table S3**). The majority of proteins that are structurally perturbed after rehydration were poorly resoluble (median resolubility 0.21), whereas those that retain native-like structure after rehydration also resolubilize to a greater extent (median resolubility 0.35). A quantile analysis of the entire dataset demonstrates a strong correlation (Pearson’s r = 0.94) between structural retention and resolubility (**Fig. 2H**). These data suggest that proteins that maintain their fold following D-R are less likely to form irreversible aggregates in the process.

### Desiccation tolerance is encoded in the surface area of folded regions

To understand the chemical and structural determinants of D-R tolerance, we first annotated proteins as membrane or non-membrane proteins, revealing 25% of proteins are annotated as being integral or peripheral membrane proteins (**Fig. 3A**). Predictably, membrane proteins are less resoluble due to their hydrophilic nature. Because the fate of membrane proteins is potentially more dependent on the lysis process (which in this case lacked detergents), our subsequent analysis focused exclusively on the 3226 non-membrane proteins.

**Fig. 3.**
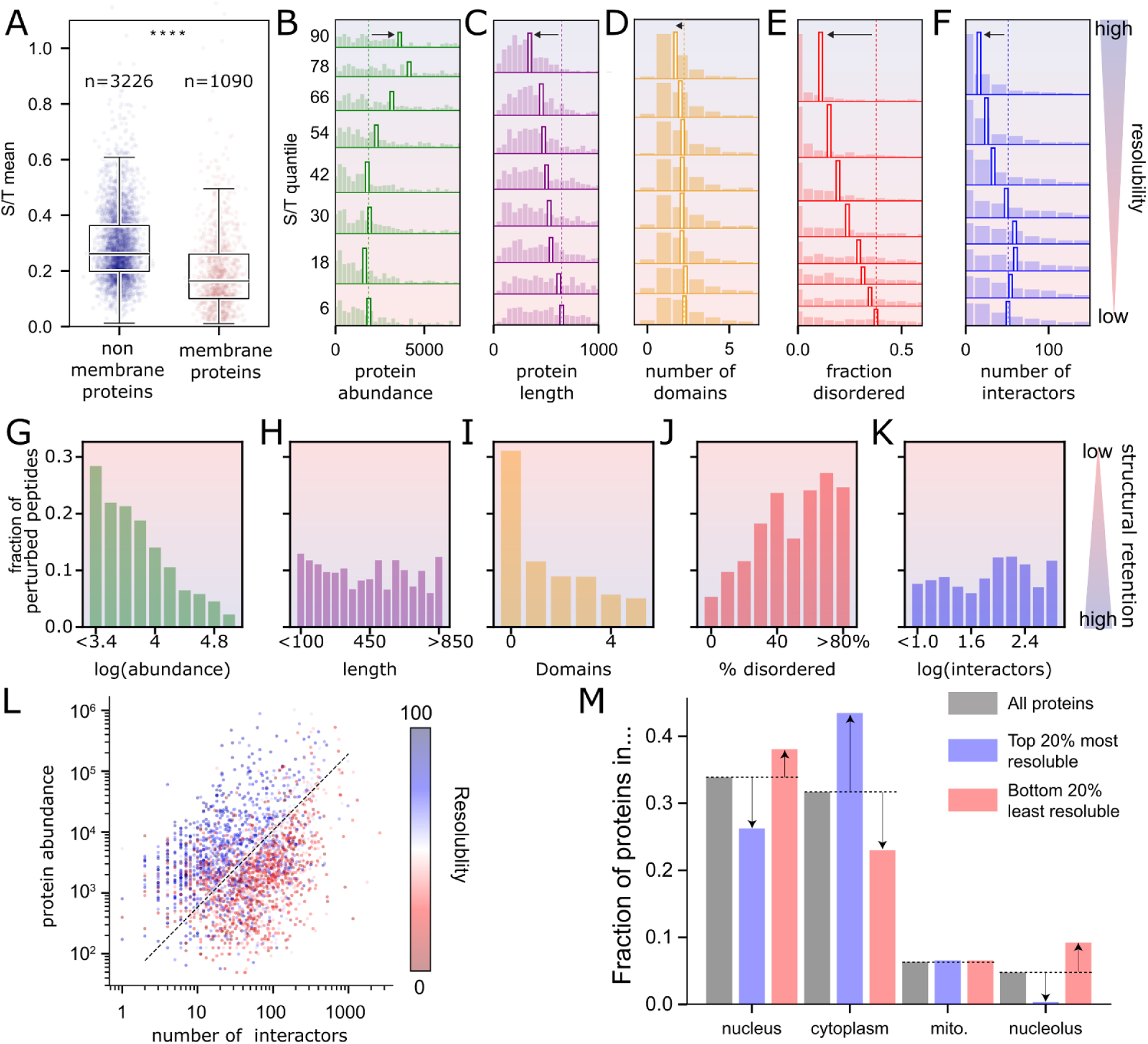
Correlates of protein resolubility and structural retention. **(A)** Within the dataset, 74% are non-membrane proteins, and 25% are membrane proteins. Membrane proteins are, on average, less resoluble than non-membrane proteins. **(B-F)** Distribution of different metrics for all non-membrane proteins in each of eight resolubility quantiles. Empty bars in each quantile represent the median, and the vertical dotted line represents the median for the lowest resolubility quantile. Distributions are shown for (B) protein abundance (in average copies per cell), (C) protein length, (D) number of folded domains, (E) percent disordered residues, and (F) the number of interactors. **(G-K)** Fraction of peptides with structural alteration (change in proteolytic susceptibility) following desiccation-rehydration as a function of their parent proteins’ (G) copy number per cell, (H) length, (I) number of globular domains, (J) fraction disorder, and (K) number of interactors. **(L)** Proteins projected into interaction count vs. copy number space, colored based on their rescaled resolubility (0 = lowest resolubility; 100 = highest resolubility). **(M)** Subcellular localization of all proteins (gray) vs. the top 20% most resoluble proteins (blue) vs. the bottom 20% least resoluble proteins (red).

We compared resolubility percentages with protein abundances (*28*). Dividing the detected, non-membrane proteins into eight S/T quantiles (**Fig. 2D**, **Table S4**), we found that resoluble proteins are generally more abundant than non-resoluble proteins (**Fig. 3B**). We also saw a systematic decrease in protein length associated with the more resoluble proteins (**Fig. 3C**) but a weak correlation with the number of distinct folded domains (**Fig. 3D**). Intriguingly, we saw a strong correlation between resolubility and disorder content (that is, the fraction of residues predicted to reside in intrinsically disordered regions). The least resoluble proteins were enriched in disordered regions, while the most resoluble proteins were depleted of disorder (**Fig. 3E**). Finally, we used STRING-DB to annotate each protein with its number of interactors in the yeast interactome (*32*). This analysis revealed that resoluble proteins tend to have fewer interactors than non-resoluble proteins (**Fig. 3F**). Overall, our analysis suggests that resolubility (i.e., D-R tolerance) is highest for yeast proteins with high cellular abundance, low disorder content, shorter length, and a low number of interactors.

For the most part, traits that make a protein more resoluble correlate with greater structure retention upon D-R (**Fig. 3G-K, Table S5**). Indeed, the most abundant proteins (>10^5^ copies/cell) are highly resistant to structural deformation with only 2.2% (86/3846) of their mapped peptides exhibiting a change in proteolytic susceptibility upon D-R, while lower abundance proteins have up to 28% of their peptides structurally altered (**Fig. 3G**). Similarly, lower disorder content is also highly correlated with structure retention (**Fig. 3J**). Moreover, we found that proteins with no annotated globular domains were greatly enriched in structural perturbations upon D-R (31% of all peptides, 153/492), supporting the view that intrinsically disordered proteins are typically altered by D-R (**Fig. 3I**). On the other hand, there was very little correlation between protein length and number of interactors with structure retention (**Fig. 3H,K**), implying that large proteins with many interactors are more likely to aggregate following D-R without structural perturbation. We found that structure retention also associates positively with an acidic isoelectric point (**Fig. S4A**) and a larger number of sites that interact with chaperone proteins (**Fig. S4B,C**).

Given how substantial of an effect disordered regions appear to have on resolubility, we extracted all the disordered regions from all the non-membrane proteins and asked whether various amino acid compositions of these regions would explain differences in resolubility. To our surprise, we could not find a correlation between any sequence features investigated across disordered regions (**Fig. S5**). This suggests that the overall sequence chemistry of disordered regions does not play a major role in determining D-R tolerance, and simply that the absence of disordered regions improves D-R tolerance while their presence undermines it.

We also examined two-dimensional spaces described by pairs of attributes and found that plotting resolubility versus both copy number (i.e., protein abundance) and interaction count creates a clear dividing surface that discriminates the most resoluble proteins as those with high copy number and few interactors and the least resoluble proteins as those with low copy number and many interactors (**Fig. 3L**). Finally, we investigated the subcellular localization of the top *versus* bottom 20% of proteins as ranked by resolubility. In general, cytoplasmic proteins tended to be more resoluble, while nuclear proteins (and nucleolar proteins especially) tended to be less resoluble (**Fig. 3M**). In consonance, cytoplasmic proteins showed high structural retention (6.6% of peptides perturbed), and nucleolar proteins were strikingly structurally altered (48.7% of peptides perturbed, **Fig. S4D**). Our analysis thus far is correlative; it illustrates attributes that tend to co-occur with D-R tolerance but fails to offer mechanistic insight into why these properties are effective. To address this, we initiated a structural bioinformatic analysis to investigate the physicochemical origins of D-R tolerance.

Given the lack of explanatory power offered by sequence features in disordered regions, and because most resoluble proteins retained their structure, we next hypothesized the core determinants of D-R tolerance may be encoded in the globular domains. Using the AlphaFold2 structural models for all *S. cerevisiae* proteins, we excised out all globular domains using two independent approaches (**Fig. 4A**) (*22*, *23*). One approach used Chainsaw, a trained, multi-track convolutional neural network for decomposing distinct subdomains (*33*). The other used DODO (https://github.com/idptools/dodo), a structure-based domain decomposition approach (**Fig. 4B**). These approaches reveal different numbers of globular domains (**Fig. S6**), and have distinct strengths and weaknesses (see *Methods*).

**Fig. 4.**
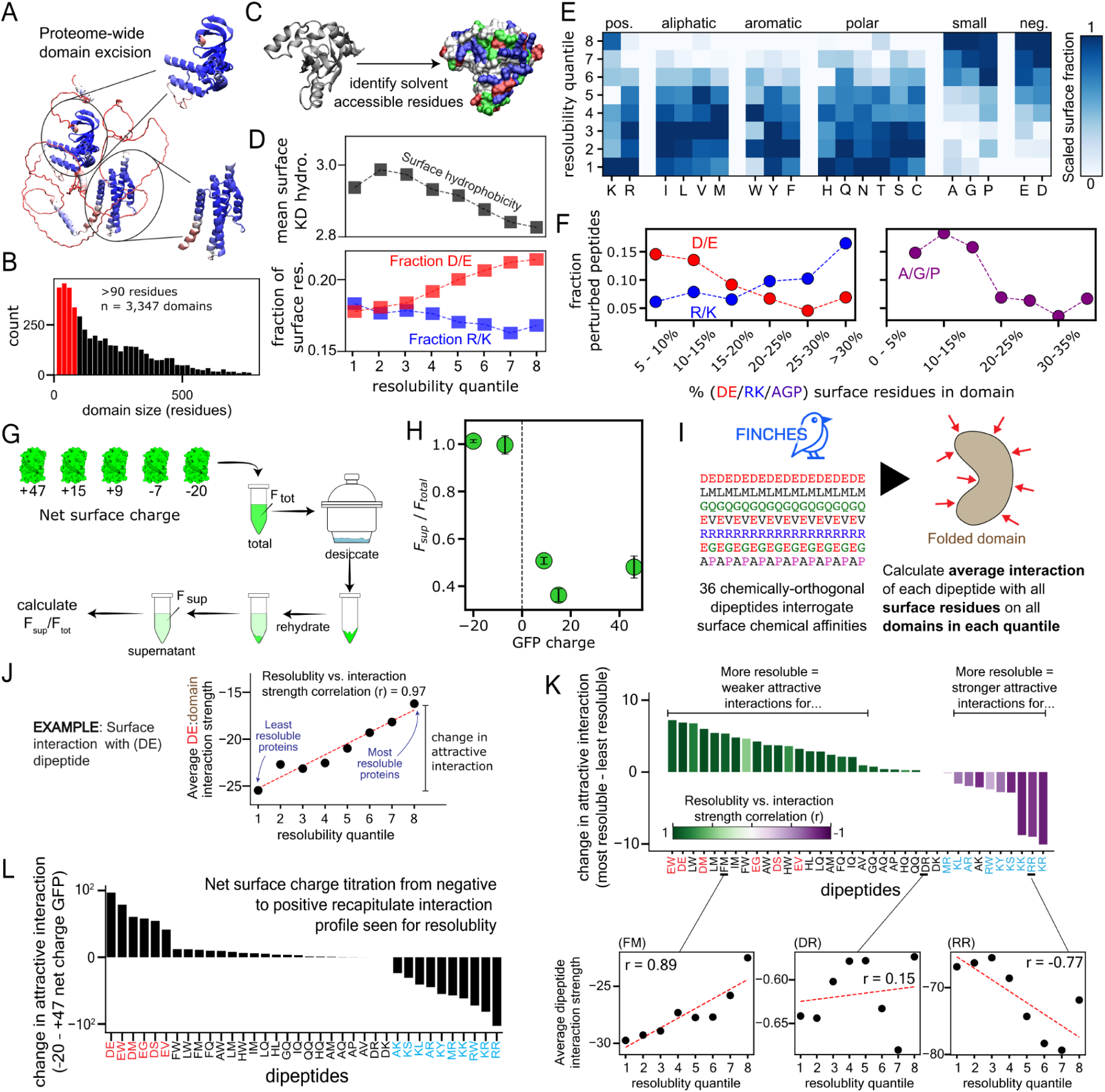
Specific surface chemistries on globular domains improve inertness and facilitate tolerance to desiccation. **(A)** Globular domains from AlphaFold2 predicted structures are excised algorithmically. **(B)** Histogram of domain size (in residues) for domain excision using DODO. Domains shorter than 90 residues (red) are not included. **(C)** Schematic representation of domain surface area quantification. **(D)** Overall hydrophobicity (top) of surface-exposed residues and fraction of surface-exposed residues that are positive (blue) or negative (red) in different resolubility quantiles. **(E)** Analysis of solvent-exposed amino acid fractions across different S/T quantiles. Each row is internally normalized such that each amino acid’s surface fraction varies between 0 and 1, enabling comparison of amino acids with very different overall abundance. (**F**) Fraction of structurally perturbed peptides from domains with increasing fraction of positive (blue), negative (red), or small amino acids (purple) at their surface. **(G)** Schematic of GFP resolubility experiment. (**H**) The ratio of GFP fluorescence after (*F_sup_*) and before (*F_total_*) D-R vs. their surface charge. (**I**) Measuring how surface amino acid compositions influence intermolecular interactions using a set of 36 chemically orthogonal dipeptide repeats. **(J)** To correlate domain interactions with resolubility, we examine the correlation coefficient of interaction vs. resolubility quantile and the magnitude of change for this interaction between the most and least resoluble quantiles. **(K)** The magnitude of change (bars) and correlation coefficients (colors) of chemical interaction vs. resolubility reveal strong chemical interaction dependencies. Note that for interaction strengths (insets) a more negative number is a more favorable interaction. **(L)** Magnitude of change of chemical interactions of dipeptide repeats between most negative and most positive GFP mutants reveals trends similar to those seen for proteome resolubilities, in line with the experimental results shown in (H).

Using individual globular domains, we identified the solvent-accessible residues (see *Methods*) and investigated the relationship between globular domain surface chemistry and resolubility (**Fig. 4C**). All results presented in **Fig. 4** use domains defined via DODO, although comparable results are obtained if the analysis is repeated using domains defined by Chainsaw (**Figs. S6-10**). We first investigated gross chemical properties, revealing that more resoluble proteins tend to have a lower surface hydrophobicity, higher surface negative charge, but curiously, lower surface positive charge (**Fig. 4D**). To obtain finer-grain insight into the role of surface chemistry, we calculated normalized surface-accessible amino acid fractions across distinct S/T quantiles in order to assess if specific residue types are more commonly found on the surface of resoluble *versus* non-resoluble proteins. Strikingly, this analysis segregated residue types into two distinct classes (**Fig. 4E**). Positively charged residues, aliphatic residues, aromatic residues, and polar residues are enriched on the domain surface of proteins with poor resolubility. In contrast, negatively charged residues and molecularly small amino acids (proline, glycine, and alanine) were all enriched on the surface of domains from highly resoluble proteins. Taken together, these results imply the surface of globular domains may be tuned to enable resolubility.

These same features also possessed explanatory power for structure retention. The LiP-MS experiment assessed the structural retention status for 1,571 domains. Domains with the lowest percentage of negatively charged surface residues showed a large fraction of peptides with perturbed structure (∼15%) compared to domains with the highest percentage of negatively charged residues (∼5%), indicating negatively charged surfaces improve structural retention upon D-R. Conversely, for domains with an abundance of positively charged residues, more structurally altered peptides were observed (>15%) compared to domains with few positive charges (5.3%) (**Fig. 4F, Table S8**). This finding echoes the patterns we found between structure retention and protein isoelectric point (**Fig. S4A**). The effect of small amino acids (Ala, Gly, and Pro) observed in resolubility is also recapitulated in structural retention: Domain surfaces depleted of these residue types are more prone to structural perturbation (14.8% of their peptides), whereas domain surfaces enriched in these residue types (≥30%) display 3.7-fold lower frequency of structural perturbation (**Fig. 4F**). Structural retention also tracked closely with resolubility for aliphatic, aromatic, and polar residues (**Fig. S7E, Table S8**), and similar trends obtained with DODO were also observed for Chainsaw domain excision (**Fig. S7F,G, Table S9**).

To test the inferences obtained from this analysis, we used a series of GFP variants with varied surface charge from +47 to –20 (**Fig. 4G**) (*34*). These variants were spiked into yeast lysates, and their fluorescence was measured before and after D-R (**Fig. 4G**). As predicted from the proteome-wide observations, these results showed negatively supercharged GFPs were almost entirely resoluble, whereas positive supercharging did not endow the protein with high resolubility (**Fig. 4H**). This result highlights the transferability of our findings to designed proteins with a high degree of desiccation tolerance through surface chemistry modifications.

Previously, we described that D-R tolerance occurs in proteins that have fewer interaction partners (**Fig. 3F,L**). Given that protein-protein interactions are typically mediated through their surfaces, we sought to understand if surface chemistry was governing resolubility by tuning interactions with other proteins. We calculated mean-field interactions between each domain’s surface residues and a set of thirty-six dipeptide repeats (see *Methods,* **Fig. 4I**) that span a diverse range of peptide chemistries, to test how different globular domain surfaces may interact with various putative partners (*35*). In total, we computed ∼500,000 domain:peptide interactions to construct a systematic map of the chemical biases encoded by every globular domain in the *S. cerevisiae* proteome.

To investigate trends between solubility and surface interactions, we averaged the interaction between each dipeptide repeat and every domain associated with proteins in each of the eight quantiles (**Fig. 4J**). This analysis allowed us to assess: (1) if there is a correlation between resolubility and average surface interaction with a given type of peptide chemistry; and (2) the magnitude of this effect; that is, how different is the interaction strength with the most (quantile 8) vs. the least (quantile 1) resoluble proteins.

This result revealed a strong and systematic trend showing that resolubility is strongly associated with weaker intermolecular interactions for most dipeptide repeats (**Figs. 4K, S8**). Strikingly, non-resoluble proteins contain globular domains that show stronger attractive interactions with hydrophobic, polar, and negatively charged test peptides. In contrast, the most resoluble proteins contain globular domains that are generally more inert: they have weak attractive interactions for those same test peptides, and show strong interactions only with the most positively charged dipeptide repeats (KK, RR, KR). This latter finding is not surprising per se, as resoluble proteins’ domain surfaces tend to be acidic, but it is worth pointing out that polycations are not a common species inside cells.

As we did for the whole yeast proteome, we also calculated how the five GFP surface-charge variants interact with the 36 test dipeptide repeats and plot in **Fig. 4L** the difference in interaction strength between the two ends of the spectrum. GFP(-20) behaved similarly to domains in the 8th quantile and GFP(+47) behaved similarly to domains in the 1st quantile, though here we observed a sharp transition and a large difference in magnitude of interactions strength between all negative and positive test peptides. As the experimental data showed a similarly sharp transition in resolubility between GFP(-7) and GFP(+9), this consistency suggests that interaction strength with negative and negative/hydrophobic peptides may be a reliable predictor for resolubility.

Based on the totality of these analyses, we conclude that proteins with amino acid surfaces that are enriched in acidic residues and residues with small sidechains are best suited for robust D-R tolerance. In contrast, surfaces with positively charged residues and hydrophobic residues tend to have poor D-R tolerance. These observations likely reflect both the intrinsic properties of certain surface chemistries being easier to solvate in the desiccated state as well as the tendency of those surfaces to be more inert to forming attractive interactions with other macromolecules.

### Tolerant proteins are producers of metabolites and small molecules

We next wanted to see if tolerant proteins have common functions. For this, we took the 20% most and least resoluble proteins (∼800 each) and examined them for functional enrichment in gene ontology (GO) and KEGG pathway association (*36–38*). Enrichment was calculated against the full set of proteins identified in the total (T) sample prior to desiccation as a reference set and significance was assigned significance based on a Fisher 1-sided t-test corrected for multiple testing using the G:SCS algorithm (*36*) (**Table S6**).

We found that the most resoluble proteins are involved in small molecule biosynthesis. A significant enrichment of amino acid and secondary metabolism synthesis was observed in KEGG, and a similar fold-change was observed for biosynthetic processes and enzymatic functions in GO (**Figs. 5A**, left). The least resistant proteins showed different functionalities, with significant enrichment of ribosome biogenesis in KEGG and related functions such as RNA processing and helicase activity in GO (**Fig. 5A**, right). This indicates that the supernatant following D-R is enriched in enzymes that process small molecules, but depleted from ribosome biogenesis machinery, arguably the greatest consumer of cellular resources (*24*). Effectively, the ability of the cell to generate new proteins could be severely dampened following D-R.

**Figure 5.**
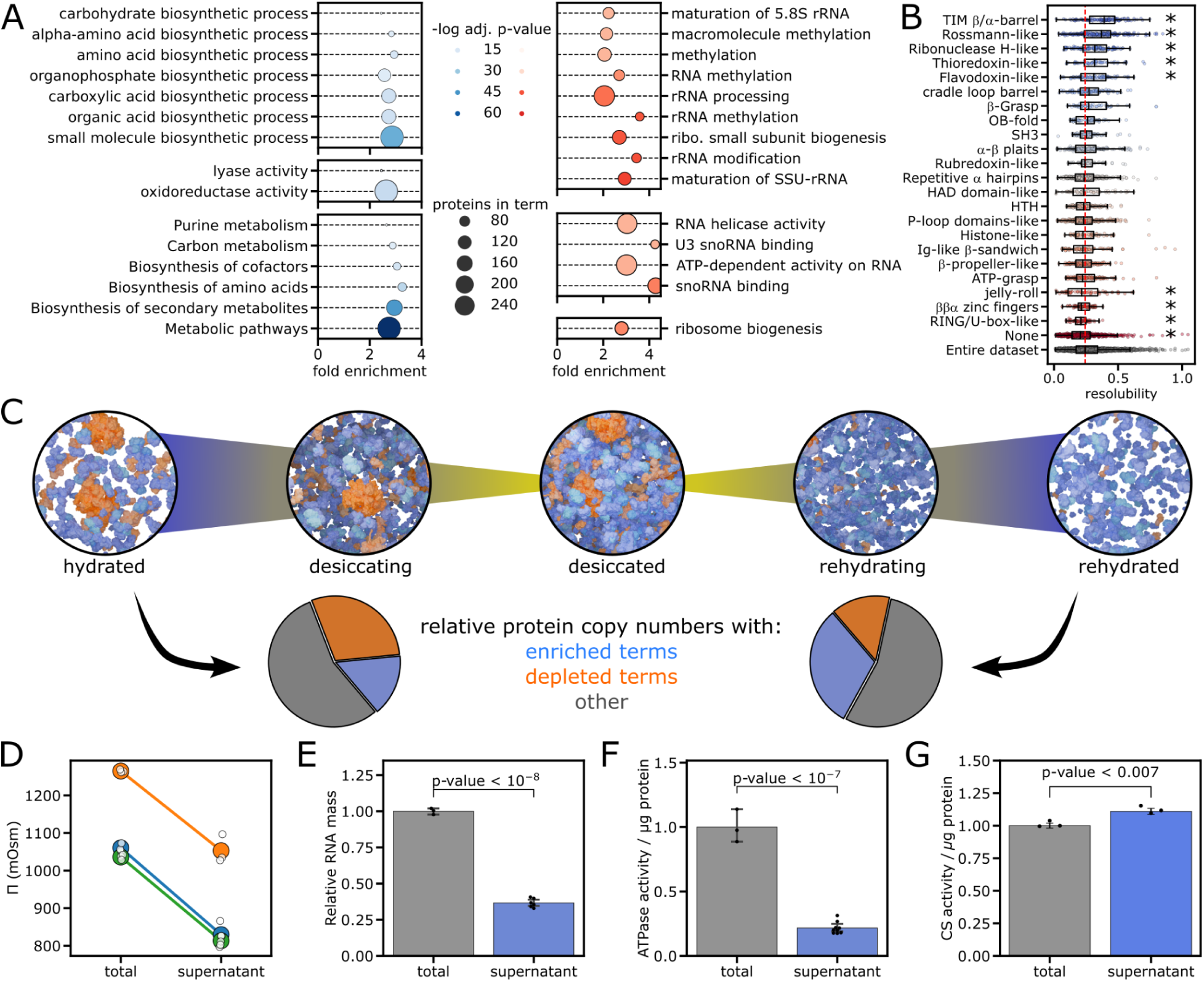
Functional enrichment following deiccation-rehydration. (**A**) Enrichment of molecular function terms in 20% most (blue) and least (red) resoluble proteins. Enrichments are for molecular processes (top), biological function (middle), and KEGG pathways (bottom). (**B**) Enrichment of ECOD domains. Asterisk denotes statistical significance (P < 0.01) by KS t-test against the entire dataset, adjusted for false-discovery. (**C**) Schematic of the metabolic kickstarting process and its effect on the protein composition of the rehydrated cytoplasm. Pie charts show lysate enrichment of GO-terms from (A) following D-R. Pie charts were calculated using copy numbers from the enriched and depleted proteins for hydrated, and the same numbers multiplied by their respective resolubilities for rehydrated charts. (**D**) Comparison of soluble lysate osmolarity before and after D-R. Colors represent different repeats of the experiment, and each point represents an average of 3 replicates. (**E**) Soluble RNA mass in the total sample before (**T**) and supernatant sample (**S**) after D-R. (**F**) ATPase activity of lysates normalized to total protein mass before (**T**) and after (**S**) D-R. (**G**) Citrate synthase activity normalized to total protein mass before (**T**) and after (**S**) D-R. In (E), (F), and (G) the points represent biological repeats taken on different days, and error bars are the standard deviation of the data. P-values are assigned via 2-sided t-test.

As an independent test of enrichment for small molecule producers, we looked for domain types (fold topologies) enriched in tolerant and non-tolerant proteins. Fold topologies are often associated with high-level biochemical functions (*39*). We assigned fold topologies using Evolutionary Classification of protein Domains (ECOD) (*40*) to each of the proteins in the dataset (**Table S7**) and identified 24 families with over 30 domains (*39*). Comparing the entire dataset, we found that TIM β/α barrels (*41*, *42*), Rossmann-like folds (*43*, *44*), Ribonuclease H-like folds, and flavodoxin folds show a significantly higher resolubility. These motifs are common in primary metabolic processes like glycolysis, electron transfer, and energy transduction (**Fig. 5B**). TIM β/α barrels, ribonuclease H-like folds, and flavodoxin folds also displayed high levels of structure retention (**Fig. S11, Table S10**). Conversely, proteins that have no assigned structural domains show significantly lower resolubility (expected, given a lack of annotated domains is correlated with intrinsic disorder (**Fig. 5B**) (*45*)). Domains associated more closely with nucleic acid binding (HTH, histone-like, and zinc fingers) typically possess lower resolubility and lower structure retention.

This analysis further suggests there is a unique functional composition in the rehydrated lysate that could be adaptive. If upon rehydration the cell resumed the typical functions of biosynthesizing ribosomes and new proteins, it would quickly be left without the molecular building blocks and metabolites required for survival. The higher tolerance of central metabolic enzymes to D-R shifts resources towards energy and metabolite production – prioritizing survival over growth (*46*, *47*) – a process we refer to as ‘metabolic kickstarting’ (**Fig. 5C**).

### Metabolic kickstarting can assist the initial stages of desiccation recovery

The ‘metabolic kickstarting’ hypothesis makes several predictions regarding metabolic activity after D-R. First, upon rehydration, the supernatant should be depleted of both small molecules and ribosomes. Second, the supernatant should be depleted of biomolecules that convert ATP to ADP; if energy consumers are less resoluble, they will be enriched in the pellet. Third, the supernatant should be enriched for biomolecules associated with cellular energy production; if energy producers are more resoluble, they will be depleted from the pellet. We tested these hypotheses directly using independent biochemical assays on lysates before and after D-R.

We first wanted to assess the depletion of small molecules following D-R. To do this, we measured the osmotic pressure of lysate before and after D-R. Our analysis revealed that there is a ca. 300 mOsm drop in osmotic pressure (**Fig. 5D**). With the proteome itself contributing only 1-5 mOsm of this (the total proteome concentration in yeast is ∼ 1-2 mM (*48*)), we conclude that the bulk of the loss results from small molecules and ions that are trapped in the desiccated pellet - by far the most abundant species in the cell(*49*). This explains the functional importance of the rehydrated proteome being enriched for small molecule producers.

Next, we tested the depletion of ribosomes following D-R. Ribosomal protein synthesis is a major energy consumer(*24*), so ribosome loss would slow down protein production and, therefore, energy consumption. To measure this, we quantified the total RNA content in our lysate before and after desiccation/rehydration. Given that ∼80% of RNA in the cell originates from ribosomal RNA, total RNA acts as a proxy for ribosome concentration (*50*). We found that while initially lysates contain a significant amount of RNA (> 8 μg/μL), there is a significant drop of 64% in the total RNA amount following desiccation and rehydration (**Fig. 5E**). This figure is qualitatively in line with the average resolubility we measured for 43 small subunit proteins (0.23 ± 0.04) and for 60 large subunit proteins (0.25 ± 0.05). Hence the supernatant following D-R appears strongly depleted of ribosomes.

To test the depletion of energy consumers, we examined the ATPase activity change in the proteome following D-R. To do this, we used a luciferase assay that measures the turnover of ATP into ADP from the lysate before D-R and in the supernatant after. This value was then normalized by protein mass, giving a value that reflects the per-protein ATP utilization. In agreement with our hypothesis, ATPase activity is reduced by nearly 75% in the supernatant, indicating that energy consumers are typically non-resoluble and/or less active in the rehydrated form (**Fig. 5F**).

Finally, to test for the enrichment of producers in the supernatant, we followed the activity of the enzyme citrate synthase (CS). Beyond being a key component of the Krebs cycle and critical for cellular energy production, CS has historically been employed as a model enzyme to assess desiccation protection (*16*, *51*). Measuring lysate CS activity before and following D-R, and normalizing this activity value per total protein mass, we find that the rehydrated lysate is significantly enriched in active CS (**Fig. 5G**). This result confirms that – at least for citrate synthase – the supernatant is enriched for functionally active energy producers. We also note that yeast CS has a somewhat high proportion of negatively charged residues on its surface (20.7%, average is 18.3%), an unusually high proportion of small residues on its surface (34.1%, average is 17%), and exhibited structural retention, suggesting that these features might also predict biochemical reactivation following D-R.

Taken together, these experiments support the metabolic kickstarting hypothesis: The rehydrated proteome is depleted of small molecules, ribosomes, and ATP consumers. On the other hand, enzymes that biosynthesize small molecules have more inert surface chemistries and generally low disorder content, which enable them to resolubilize more efficiently following D-R, thereby helping yeast survive this severe stress.

## Discussion

The tolerance of the proteome to desiccation events is crucial for understanding cellular resilience and survival under extreme conditions. By examining the proteome of a well-established model organism in the absence of protection afforded by priming, we aimed to uncover the basal tolerance encoded in protein sequence and structure following desiccation-rehydration (D-R). The neat lysates we produced for this work resemble the composition, pH, and solute concentration conditions encountered in the specific state the cells were in when lysed. This allows us to observe protein behavior in a complex environment that is more physiologically relevant than afforded by most *in vitro* studies.

We specifically analyzed the proteome of yeast in the logarithmic phase, in which cells do not exhibit desiccation resistance because they have not upregulated chaperones or produced protective osmolytes (*25*). This intentional choice provides a baseline for understanding the intrinsic tolerance of each protein when exposed to desiccation in the context of the cellular environment.

Our results revealed that approximately 35% of the proteins remain soluble after the D-R cycle. While this finding aligns with expectations (*52*), proteome-wide analysis deconstructs this result to reveal a complex landscape of desiccation resistance across the *S. cerevisiae* proteome. Nearly all proteins were present at detectable amounts in the rehydrated supernatant. This indicates that even poorly resistant proteins have some copies solvated following rehydration. However, resolubility is not sufficient: to function, folded proteins must also retain their native states. To test this, we used limited proteolysis (LiP) to profile protein structures. Our results show a high degree of correlation between resolubility and structural retention following D-R. This correlation is observed here for the first time on a proteomic scale, and suggests aggregation and loss of native structure are well correlated. Nonetheless, these two measures sometimes differ. For instance, proteins with many interactors are not more likely to be structurally perturbed themselves, but they generally exhibit low resolubility, possibly because they bind to *other* proteins that become structurally perturbed and aggregate.

The most resistant proteins are small, bereft of disordered regions, and abundant. They also tend to have smaller interactomes and less attractive interactions with many “test” peptides based on computational predictions. This indicates that resolubility is a more complex feature than mere solubility: a lack of strongly interacting surfaces is important to help proteins escape the complex matrix of a desiccated cytoplasm. We were surprised to find that disordered regions are actually counterproductive for D-R resistance - in marked contrast to what was observed for their role in rescuing proteins from denaturant-induced unfolding (*21*). However, desiccation places the proteome in a profoundly different state than heat shock or denaturant (*14*, *19*). In the desiccated state – where proteins become ultra-concentrated – the higher potential for disordered regions to interact with other species may render them a liability.

One key factor that governs resolubility is the average surface chemistry of folded domains. Globular domain surface composition has already been shown to correlate with stress resistance on a few purified proteins (*34*), as well as similar observations in extremophiles (*53*, *54*). Here, we show that increasing negative charges alone is enough to dramatically enhance the resolubility of fluorescent proteins in rehydrated lysates. This finding suggests that proteins can be made resistant to desiccation by modifying their surface charges. This observation is logical: surface-exposed residues are subject to higher evolutionary rates than residues in a protein’s hydrophobic core (*55*, *56*), and for monomeric enzymes with small interactomes, they can be tuned with relatively few restrictions. This observation also aligns with the bioengineering practice of “supercharging” proteins by mutating many surface residues, which has been found to improve protein stability in a number of cases (*34*, *57*). It is also reasonable that the most abundant proteins have a surface area that is less prone to desiccation: high-abundance proteins were found to exist on the edge of solubility even in well-hydrated environments (*1*), and a shift in solubility was found to occur in cellular aging (*2*), creating a constitutive pressure to make such proteins less susceptible to aggregation.

We speculate the bias for negative charge over positive charge reflects the nature of the counterion these surface residues will likely embrace in the desiccated state. The most abundant cation in yeast is K^+^, whereas the most abundant anion is glutamate (*56*, *58*). In the desiccated form, negative surfaces will become sheathed with K^+^ that easily repartition into solution during rehydration. Positive surfaces would become coated with glutamate, fatty acids, nucleic acids, and other potentially “stickier” interactions that are more challenging to resolubilize. In line with this, recent work suggests that polycations are inherently toxic to cells, and essentially every abundant protein with a strong net positive charge is neutralized by a constitutive counterion (*59*). Put succinctly, if resolubility requires a biologically inert surface, minimizing interactions with polyanions (e.g., phospholipids, nucleic acids, microtubules, *etc.*) or other proteins is enabled by enriching the surface of negative charge and depleting it of hydrophobicity and positive charge. We hypothesize the chemical determinants of resolubility uncovered here should be broadly applicable across the kingdoms of life.

Perhaps most surprising is the correlation between protein function and their desiccation survival. We find that small-molecule-producing enzymes are highly enriched in the rehydrated supernatant, while proteins associated with ribosome biogenesis are highly enriched in the least resistant fraction. Depleting the rehydrated supernatant of ribosomes reduces the synthesis of new protein machinery, allowing the cell to focus on rebuilding its reserve of building blocks, and slowly recuperate metabolism following rehydration. As desiccation is an environmentally relevant stressor yeast must face, we propose that abundant enzymes responsible for small molecule production have evolved to survive D-R by adapting their surface chemistry to increase tolerance. It is also possible that resistance to D-R provided negative selective pressure that prevented metabolic enzymes from acquiring disordered regions.

Overall, our results point to a molecular mechanism, rooted in protein surface chemistry, that helps a specific functional subset of proteins survive desiccation and rehydration - a critical stressor in many ecosystems. How the introduction of protective protective molecules such as osmolytes and chaperones may affect D-R resistance remains a question of ongoing interest (*60*, *61*). Finally, these findings suggest that protein surfaces can be rationally engineered to facilitate survival from desiccation stress.

## Data and Materials Availability

The raw mass spectrometry data used in this work can be obtained from PRiDE (Project PXD055855 [Reviewer access Username; Password: z094Yr2wIkm5] and PXD055914 [Reviewer access Username; Password: 2Cxq0Zou4cgj]).

All code used to produce figures and perform analyses can be found at https://github.com/sukeniklab/Romero_Surface_Desiccation.

## Acknowledgements

Support for this project came from an NSF Collaborative Proposal via the Integrative Research in Biology (IntBIO) program under awards 2128069 to TCB, 2128068 to ASH, and 2128067 to SS. SS and SDF thank the support from the Sloan Foundation. A.S.H. was supported by the Human Frontiers in Science Program (RGP0015/2022). JML was supported by the National Science Foundation via grant number DGE-2139839. HMM thanks the Chemistry-Biology Interface Program training grant (NIH-T32GM080189-13). EMS thanks the Program in Molecular Biophysics training grant (NIH-T32GM135131). SDF thanks the NSF Division of Molecular and Cellular Biology for a CAREER grant (MCB-2045844) as well as support from a Camille Dreyfus Teacher-Scholar Award. SS, TCB, HL, and JI are supported by The Water and Life Interface Institute (WALII), NSF DBI grant #2213983, and members of WALII provided helpful discussions. We thank the “Extreme Biophysics - The Molecular Limits of Life” research coordination Network (NSF award 1817845) in which some of the collaborations were established. Special thanks to Bruker Proteomics Applications Scientists Matthew Willetts and Diego Assis for their crucial assistance with LC-MS Data Acquisition and analysis.

## Materials and Methods

### Saccharomyces cerevisiae culture

A single colony of *Saccharomyces cerevisiae* strain BY4742 (S288C) (*62*), picked from a fresh YPD plate, was used to inoculate 250 mL of YPD medium. Yeast was grown at 30 °C and 200 rpm overnight. The overnight culture was then used to inoculate 2 X 2L flasks containing 1 L of YPD medium. Cultures were initiated with an optical density at 600 nm (OD_600_) of 0.07 and collected by centrifugation when the OD_600_ reached the mid-log phase (0.6).

### Neat lysate preparation

Yeast cells grown to mid-log phase, were resuspended in 2 mL of water, and subsequently vacuum filtered on a 1.2 μm pore cellulose membrane to eliminate media residues. Approximately 4 g of cells were obtained from a 2 L log-phase yeast culture. The filtered cells were promptly submerged in liquid nitrogen and then lysed via mechanical grinding using a Spex 6850 Freezer Mill. The cryogrinding protocol involved 4 minutes of rest followed by 9 cycles of 1 minute of grinding at a rate of 9 Hz and 1 minute of rest. A measured mass of the resulting fine powder was then transferred to a fresh 50 mL falcon tube containing Protease Inhibitors (PI). The required volume of PI 100x stock solutions (0.5 mM PMSF, 0.015 mM E-64, and 0.05 mM Bestatin in DMSO) was determined using a yeast cell density of 1.15 g/mL as the reference point. The ground cells were allowed to thaw with gentle shaking at 4°C. Following thawing, cell debris was removed by centrifugation at 3,220 x g for 20 minutes in a refrigerated tabletop centrifuge set at 4°C. The resulting supernatant was then subjected to a second spin at 16,000 x g for 15 minutes at 4°C. The supernatant, referred to as the neat-lysate or total (T) fraction, was carefully transferred to a fresh tube. The protein concentration in the neat-lysates was determined using the bicinchoninic acid assay (Pierce™ Rapid Gold BCA Protein Assay Kit, Thermo Scientific).

### Desiccation-rehydration treatment

A microfuge tube holding an initial volume of 0.1 mL of neat lysate was vacuum-dried in a desiccator for ∼30 hours until it attained a stable weight. The rehydration process involved adding water to double the sample’s initial mass. The rehydrated sample was gently agitated on a rotisserie at 4°C for 10 minutes, subjected to pipetting up and down for 5 minutes and subsequently re-incubated at 4°C for 1 hour using the rotisserie. The resoluble fraction, also known as the Supernatant (S), was obtained by centrifuging the rehydrated sample at 16,000xg for 15 minutes at 4°C. The supernatant fraction was then meticulously collected, and its volume was recorded to ensure accurate mass balance calculations. The remaining Pellet (P) fraction was resuspended in 1 mL of 8M urea solution. Subsequently, the protein concentration in both the Supernatant (S) and Pellet (P) fractions was determined using the bicinchoninic acid assay (BCA Protein Assay Kit, Thermo Scientific).

### Mass balance

The volume and concentration of each fraction were documented to evaluate the conservation of protein mass. Utilizing this information, we estimated the distribution of proteins between the S and P fractions and employed partitioning factors to scale the individual protein intensities recorded by MS.

### Thermogravimetric analysis

Samples were run on a TA TGA5500 instrument in 0.1 mL platinum crucibles (TA 952108.906). Crucibles were tared before sample loading. Crucibles were loaded with between 5 mg and 10 mg of sample. Each sample was heated from 30 °C to 250 °C at a 10 °C per minute ramp.

Determination of water loss was conducted using TA’s Trios Software (TA Instruments TRIOS version # 5.0.0.44608). The Trios software “Smart Analysis” tool was used to identify the inflection points along the Weight (%) curve and the 1st derivative Weight (%) curve. The Trios software “Step transition” tool was then used to select the area of the Weight (%) curve from the start of the run to the rightmost region of the curve that corresponded with the last flat region of the 1st derivative Weight (%) curve. The Trios software “Onset” and “Endset” tools were then used to identify the onset and endset of water loss.

### Lysate characterization

The osmolality of neat lysates was measured using a Wescor VAPRO 5520 (Wescor, Logan, UT), calibrated with manufacturer-provided standard solutions of known osmotic potentials (100, 290, and 1000 mmol/kg). For each trial, 10 μL of the total fraction and 10 ul of the supernatant fraction lysate were placed on a filter paper disk and measured. Triplicate measurements were taken for each sample. The pH was measured in triplicate for each of the total and supernatant fractions. For each measurement, 50 μL of the sample was used. A Mettler Toledo InLAB Micro pH probe attached to a Mettler Toledo SevenCompact Duo pH/Conductivity meter were employed for the pH measurements

### GFP Spike Experiments

Neat lysates were spiked with green fluorescence protein (-30 GFP, -20 mEGFP, +9 GFP, +15 GFP, +47 GFP) to a final 1 μM. To assess the active concentration of the protein, Fluorescence was measured using λ = 488 ± 25 nm excitation light, and read between 490 and 600 nm emission using a BMG LabTech ClarioSTAR. Fluorescence values before and after D-R were measured in triplicates from the spiked total sample and after D-R from the supernatant sample. The total fluorescence is reported as the ratio between the supernatant and total, and taken from the average emission between 505 nm to 515 nm. Errors are standard deviations of the average between triplicates.

### Proteome-wide Resolubility Sample Preparation

200 μg of protein from the Total (**T**), Supernatant (**S**), and Pellet (**P**) fractions were diluted in 200 μL of Native Buffer (20 mM HEPES-KOH pH 7.4, 2 mM MgCl2). Samples were then transferred to fresh microfuge tubes containing 152 mg urea. Next, 4.5 μL of a freshly prepared 700 mM stock of DTT was added to each sample, and the mixture was incubated at 37 °C for 30 minutes at 700 rpm on a thermomixer to reduce cysteine residues. Following reduction, an 18 μL portion of a freshly prepared 700 mM stock of iodoacetamide (IAA) was added to each sample, and the mixture was then incubated at room temperature in the dark for 45 minutes to alkylate the reduced cysteine residues. After alkylation of the cysteines, 942 μL of 100 mM ammonium bicarbonate was added to each sample to dilute the urea to a final concentration of 2 M. Subsequently, a 1 μL portion of a 1 mg/mL stock of trypsin (NEB Trypsin-ultra^TM^, Mass Spectrometry Grade) was added to each sample and the mixtures were incubated overnight at 25 °C at 700 rpm.

Sep-Pak C18 1cc Vac Cartridges (Waters) were employed using a vacuum manifold to desalt the peptides. The clean-up procedure, outlined in (*63*) began with acidification of the peptides using 12.6 μL of trifluoroacetic acid (TFA). Subsequently, the cartridges were conditioned with 1 mL of Buffer B (comprising 80% ACN and 0.5% TFA), and equilibrated with 4 × 1 mL of Buffer A (0.5% TFA). Peptides were then loaded onto the cartridges and subjected to a wash step using 4 × 1 mL of Buffer A. Elution of peptides was achieved by adding 1 mL of Buffer B into the columns. Vacuum cartridges were placed inside 15 mL conical tubes and centrifuged at 350 g for 8 minutes. The eluted peptides were transferred to microfuge tubes and vacuum-dried before being stored at −80 °C until MS analysis.

### Proteome-wide Resolubility LC-MS Data Acquisition

LC–MS was performed on a NanoElute (Bruker Daltonik) system coupled online to a hybrid TIMS-quadrupole TOF mass spectrometer (*64*) (Bruker Daltonik timsTOF Pro2, Germany) via a nano-electrospray ion source (Bruker Daltonik Captive Spray). Approximately 200 ng of peptides were separated on an Aurora column 25 cm × 75 µm ID, 1.7 µm reversed-phase column (Ion Opticks) at a flow rate of 250 nL min-1 in an oven compartment heated to 50 °C. To analyze samples from whole-proteome digests, we used a gradient starting with a linear increase from 5% B to 25% B over 25 min, followed by further linear increases to 37% B in 4 min and to 90% B in 4min, which was held constant for 5 min. The column was equilibrated using 4 volumes of solvent A.

The timsTOF was operated in dia-PASEF mode (*65*) with mass range from 100 to 1700 m/z, 1/k_0_ Start 0.75 V s/cm^2^, End 1.30 V s/cm^2^, ramp and accumulation times set to 50 ms, Capillary Voltage was 1600 V, dry gas 3 L/min, and dry temp 200 °C.

dia-PASEF settings were set to an optimized acquisition scheme using py_diAID software (*66*) with dia-windows isolation starting from 7 to 95 Da according to the multiple charge ion cloud density map. Each cycle consisted of 1x MS1 full scan and 60x MS2 windows covering 350 – 1250 m/z and 0.7 - 1.30 1/k_0_ (see supplementary material). The cycle time was 1.12 s. CID collision energy was 20 eV (0.60 1/k_0_) to 59 eV (1.6 1/k_0_) as a function of the inverse mobility of the precursor.

### Proteome-wide Resolubility LC-MS Data Analysis

The Spectronaut18 proteomics analyzer (*67*) was utilized with the directDIA+ (Deep) workflow to analyze spectra and perform Label-Free Quantification (LFQ) of detected peptides with a 1% false-discovery rate (FDR). A tryptic search allowing up to 2 missed cleavages for peptides ranging 7-52 residues was conducted against the *S. cerevisiae* (UP000002311, UniProt) reference proteome. Methionine oxidation and N-terminus acetylation were allowed as dynamic modifications. Carbamidomethylation of cysteines was specified as a static modification. All ratios reported here (S/T, S/P, and S/S+P) were determined from protein-level intensity measurements (as calculated from areas under the peaks) averaged across three replicates, which were then normalized by protein mass as determined by BCA assay. The p-value is calculated from the standard deviation between replicates, and corrected for FDR.

### Limited Proteolysis Mass Spectrometry (LiP-MS) Sample Preparation

The LiP assay was conducted according to the procedure outlined in the referenced literature (*68*). In summary, 2 µL of a PK stock solution (0.1 mg/mL PK in a 1:1 mixture of Native Buffer and 20% glycerol) was dispensed into a sterile 1.5 mL microcentrifuge tube. Total (T) and Supernatant (S) samples, each containing 200 μg of protein diluted in 200 μL of Native Buffer, were prepared.. To initiate LiP, the samples were combined with the PK-containing microcentrifuge tube and vigorously mixed by rapid vortexing, followed by immediate centrifugation to sediment the liquids at the bottom of the tube. Samples were incubated for exactly 1 min at 25°C. Afterwards, the microfuge tubes containing the LiP samples were transferred to a mineral oil bath pre-equilibrated at 105°C for 5 min to quench PK activity. Boiled samples were then flash centrifuged. LiP samples were transferred to 152 mg of urea. Next, 4.5 μL of a freshly prepared 700 mM stock of DTT was added to each sample, the mixture was incubated at 37 °C for 30 minutes at 700 rpm on a thermomixer to reduce cysteine residues. Following reduction, an 18 μL portion of a freshly prepared 700 mM stock of iodoacetamide (IAA) was added to each sample, and the mixture was then incubated at room temperature in the dark for 45 minutes to alkylate the reduced cysteine residues. After alkylation of the cysteines, 942 μL of 100 mM ammonium bicarbonate was added to each sample to dilute the urea. Subsequently, a 1 μL portion of a 1 mg/mL stock of trypsin (New England Biolabs, Inc. Typsin-ultraTM, Mass Spectrometry Grade) was added to each sample, and the mixtures were incubated overnight at 25 °C at 700 rpm. The following day, peptides were cleaned up using Sep-Pak C18 1cc Vac Cartridges (Waters) and stored as discussed in the <u>Proteome-wide</u> <u>Resolubility Sample Preparation methods section.</u>

### LiP-MS Data Acquisition

Chromatographic separation of digests were carried out on a Thermo UltiMate3000 UHPLC system with an Acclaim Pepmap RSLC, C18, 75 μm × 25 cm, 2 μm, 100 Å column. Approximately, 1 μg of protein was injected onto the column, which was maintained at 40 °C. The flow rate was 0.300 μL min-1 for the duration of the run with Solvent A (0.1% FA) and Solvent B (0.1% FA in ACN) used as the chromatography solvents. Peptides were allowed to accumulate onto the trap column (Acclaim PepMap 100, C18, 75 μm x 2 cm, 3 μm, 100 Å column) for 10 min (during which the column was held at 2% Solvent B). The peptides were resolved by switching the trap column to be in-line with the separating column, quickly increasing the gradient to 5% B over 5 min and then applying a 95 min linear gradient from 5% B to 40% B. Subsequently, the gradient held at 40% B for 5 min and then increased again from 40% B to 90% B over 5 min. The column was then cleaned with a sawtooth gradient to purge residual peptides between runs in a sequence. A Thermo Q-Exactive HF-X Orbitrap mass spectrometer was used to analyze protein digests. A full MS scan in positive ion mode was conducted and followed by 20 data-dependent MS scans. The full MS scan was collected using a resolution of 120000 (@ m/z 200), an AGC target of 3E6, a maximum injection time of 64 ms, and a scan range from 350 to 1500 m/z. The data-dependent scans were collected with a resolution of 15000 (@ m/z 200), an AGC target of 1E5, a minimum AGC target of 8E3, a maximum injection time of 55 ms, and an isolation window of 1.4 m/z units. To dissociate precursors prior to their reanalysis by MS2, peptides were subjected to an HCD of 28% normalized collision energies. Fragments with charges of 1, 6, 7, or higher and unassigned were excluded from analysis, and a dynamic exclusion window of 30.0 s was used for the data dependent scans.

### LiP-MS Data Analysis

The FragPipe v20.0 proteomics pipeline with IonQuant v1.9.8 with a match between runs (MBR) false discovery rate (FDR) of 5% was utilized to analyze spectra and perform Label-Free Quantification (LFQ) of detected peptides (*69*, *70*). Using MSFragger v3.8 (*71*) and Philosopher v5.0 (*72*), a semitryptic search allowing up to 2 missed cleavages was conducted against the *S. cerevisiae* (UP00000231, UniProt) reference proteome, and identifications were filtered to a 1% FDR. An MS1 precursor mass tolerance of 10 ppm and an MS2 fragmentation tolerance of 20 ppm was used. Methionine oxidation and N-terminus acetylation were allowed as dynamic modifications, while carbamidomethylation on cysteines was defined as a static modification. Raw ion intensity data for identified peptides were exported and processed utilizing FLiPPR (v0.1.4) (*31*). Data were merged from the ion to the peptide level (**Table S4**) and normalized using the log2 (S/T Protein Abundance Ratio) as outlined in the resolubility study. In all cases, missing data imputation, filtering, and Benjamini-Hochberg FDR correction were carried out per-protein, as implemented in FLiPPR. In proteome-wide analyses, peptides were labeled significantly “structurally perturbed” by desiccation-rehydration if their normalized abundance changed by more than 2-fold between S and T samples and adjusted P-values were less than 0.05. Peptides were normalized by the parental proteins’ resolubility value (average S/T) if the resolubility was less than 50% (i.e., >2-fold effect size) and the P-value was < 0.01. To label a protein or domain “structurally perturbed,” it needed to have two or more structurally perturbed peptides mapped to it. To label a protein or domain “structurally retained,” it needed to have no more than one structurally perturbed peptide and two or more total peptides assigned to it.

### ATPase activity assays

ATPase activity in the Total and Supernatant fractions was assessed using the ATPase Assay Kit (abcam ab270551), which detects inorganic phosphate (P_i_) colorimetrically as a direct product of ATPase activity. Given the intrinsic presence of ATP and P_i_ in neat lysates, the T and S fractions were diluted with Native Buffer to a concentration of approximately 2 μg/μL. Subsequently, 100 μL of the diluted fractions were processed through Zeba™ Spin Desalting Columns with a 7K MWCO to eliminate small solutes, including P_i_ and ATP. The ATPase assay followed the kit instructions, utilizing protein quantities ranging from 0.25 to 5 μg from each fraction, and was performed at 30°C for 1 hour. We prepared assay blanks without protein from either fraction to serve as controls. The signal from these blanks was subtracted from the samples’ signals to ensure accurate measurements. Results were expressed as relative ATPase Activity, calculated by dividing the absorbance at 624 nm by the mass in μg of protein in the Supernatant fraction and then further dividing by the analogous value of the Total fraction. This ratio represents the fold change or percentage relative to the Total sample.

### Citrate Synthase Activity

The Citrate Synthase Activity in the Total and Supernatant fractions was evaluated using the Citrate Synthase Activity Assay Kit (Sigma-Aldrich MAK193). This assay detects -SH groups produced during the enzymatic conversion of oxaloacetate and acetyl-CoA to citrate by citrate synthase. The reaction progress was monitored at 412 nm over 90 minutes. Following the kit instructions, we measured citrate synthase activity using protein quantities ranging from 100 to 400 μg from each fraction in triplicates. Due to significant background levels in the samples, a sample blank was included for each sample by omitting the substrate from the reaction mix. The readings from the blanks were subtracted from all sample readings to account for background interference and ensure accurate measurement of the enzymatic activity.

The results were expressed as Relative Citrate Synthase Activity by dividing the change in absorbance at 412 nm per minute per microgram of protein in the Supernatant fraction, and then further dividing by the analogous value of the Total fraction. This ratio represents the fold change or percentage relative to the Total sample and was averaged between the triplicates. The errors are standard deviations of the average between triplicates.

### RNA purification and Quantification

4-20 uL of T and S fraction was used for RNA purification. The RNeasy Plus Mini Kit was used following the manufacturer’s procedure. RNA quantification was done using the Qubit™ RNA Broad Range (BR) kit. Measurements were taken in triplicates and averaged. Errors are standard deviations of the averages.

### Protein-based Analyses

Proteins, along with their mean S/T (resolubility) value, were paired with various metadata compiled from various sources, including the Saccharomyces Genome Database (*73*), ECOD (*39*), and DomainMapper (*45*). Protein abundance data was obtained from Ghaemmaghami and SGD, as described previously (*28*, *74*). Protein length was based on the total number of amino acids in each protein sequence. The fraction disordered is calculated as the fraction of amino acids in each protein that falls within a contiguous disordered region, as identified by metapredict V2-FF. The number of interactors was determined by taking physical interaction partners reported in the STRING database (specifically, we used the data from **`4932.protein.physical.links.v12.0.txt`**) and mapping interprotein interaction partners (*32*). Subcellular localization annotations were taken from UniProt, and only the top four most commonly observed annotations were highlighted.

Membrane proteins vs. non-membrane proteins were segmented using UniProt annotations based on subcellular localization. Annotations were obtained on June 14th, 2024. This identified 1775 membrane proteins and 4264 non-membrane proteins across the yeast proteome. After cross-referencing with proteins recovered in our mass spectrometry experiments, this left 1090 membrane proteins and 3226 non-membrane proteins. Membrane proteins here include both integral and peripheral membrane proteins.

Statistical analyses of resolubility were based on dividing the 3226 quantified non-membrane proteins into eight resolubility quantiles based on ranking their S/T and assessing the distribution of protein properties of interest within that quantile. The set of proteins in each quantile is given in **Table S3**.

Statistical analyses of structure retention were based on dividing proteins by properties of interest, and then counting the number of structurally perturbed and non-perturbed peptides associated with all proteins in that category. Proteins with their associated metadata and peptide counts are given in **Table S5**. We note that structure retention analyses did ***not*** explicitly strip out membrane proteins, because LiP-MS provides local structurally-resolved information, and can provide valid information about soluble/globular portions of membrane-localized proteins.

### Structural Domain-based Analyses

Proteome-wide structural bioinformatics was based on the AlphaFold2 *S. cerevisiae* proteome (UP000002485) obtained from https://alphafold.ebi.ac.uk/download#proteomes-section in March 2024 (*22*, *23*). This initial dataset contained 6039 yeast proteins. From those proteins, we excised out individual intrinsically disordered regions using metapredict (V2-FF) for disorder analysis (see below) (*75*, *76*). Disordered regions were identified using the `predict_disorder_domains()` function in metapredict (V2-FF). All informatics used SHEPHARD to organize, parse, and sanity-check protein sequence information and annotations, while all sequence-based analysis was performed using sparrow or FINCHES (*35*, *74*) (https://github.com/idptools/sparrow). All code for the structural domain-based analyses are at https://github.com/holehouse-lab/supportingdata/tree/master/2024/romero_2024.

We also excised out globular folded domains using two distinct bioinformatic approaches. One approach used chainsaw, a supervised learning approach trained to predict whether a pair of residues comes from the same contiguous domain (*33*). The other used DODO, a structural informatics approach that approximates folded domain boundaries by quantifying the number of atoms within a specific threshold distance of all other atoms in a given structure. For chainsaw, we performed domain segmentation on the structure predictions of the S. cerevisiae proteome obtained from the AlphaFold Database. Chainsaw predictions were performed with default parameters - e.g., `--remove_disordered_domain_threshold` left at 0.35 and `--min_domain_length` was left at 30 residues and `--min_ss_components` left at 2. For DODO, default settings were used with the `build.pdb_from_pdb(input_file, output_file, just_fds=True)` command.

Broadly speaking, two types of error can occur in the context of domain decomposition. One type occurs when an algorithm is overly aggressive, segmenting domains that should not be segmented and creating artificial interfaces that incorrectly expose residues. Another type occurs if an algorithm is insufficiently sensitive, grouping multiple domains together into a single large domain. These two types of errors have distinct impacts on the types of residues reported as solvent accessible. We, therefore, sought to apply two different approaches for domain decomposition, each of which were biased towards opposing error types. Thus, if both approaches led to the same conclusion, this would confirm the conclusions were robust to the details and biases of the domain decomposition approach.

Chainsaw was used in a mode whereby discontinuous domains were segmented apart, effectively ensuring that it would behave in an overly aggressive way. In contrast, given DODO was developed for the reconstruction of disordered regions around folded domains, as opposed to domain decomposition, DODO often erred on the side of failing to decompose multi-domain subunits into distinct globular domains. We therefore decomposed the yeast proteome and performed all analyses discussed here in these two ways, with the goal of providing confidence that any conclusions drawn were not due to idiosyncrasies of a specific algorithmic approach to domain decomposition.

Resolubility analysis of structural domains focused on non-membrane proteins that have at least one domain between 90 and 1000 amino acids in length. This filtering leaves us with 3347 domains (DODO) or 5477 domains (chainsaw) spread over 2667 proteins (DODO) or 2884 proteins (chainsaw).

Solvent-accessible residues were identified using individual globular domains with FINCHES, which in turn uses the Shrake Rupley algorithm implemented within MDTraj (*35*, *77*). Solvent accessibility of each residue was determined in A^2^ using a probe radius of 0.14 nm on a per-residue basis. Whether a residue was solvent accessible or not was defined as to whether the total residue SASA was over 40% accessible of the maximum possible accessibility for that residue type. Maximum accessibility was determined by performing all-atom simulations of GXG tripeptides and computing the average accessibility of the guest peptide. Various accessibility percentages were compared, and 40% offered a reasonable tradeoff between being sufficiently permissive to capture sidechain conformers that may occlude their accessibility in the specific AlphaFold2 structure being used and being sufficiently restrictive to prevent residues deep in crevices from being included. We also repeated our entire pipeline using a 10% threshold and found no difference in any of the conclusions; consequently, the trends are robust to the specific details regarding solvent accessibility. **Fig. S9** compares the fraction of residues that are solvent accessible vs. domain size (in amino acids).

The solvent-accessible amino acid composition was computed in two different ways: by the overall surface area of a given residue and by the overall count of a given residue. In both cases, the overall approach involved taking each individual DODO (or chainsaw) domain, identifying solvent-accessible residues, and then either counting the number of each residue type that was solvent accessible or taking the summed surface area for each different residue type. These values were added to a per-residue tally, which was updated for each domain. For surface hydrophobicity, the average Kyte-Doolitle value of all surface-exposed residues was computed (**Fig. 4E**). The fraction of surface residues that are positive (Arg/Lys), negative (Asp/Glu), small (Ala/Gly/Pro), aromatic (Phe/Tyr/Trp), hydrophobic (Ile/Leu/Val/Met), or polar (all others)was computed (**Fig. 4E**). For individual amino acids, the fraction of surface residues made of each amino acid was calculated (**Fig. 4F**). We also calculated the fraction of the overall globular domain surface area made up of each amino acid (**Fig. S10**). In both cases, the same overarching conclusions emerge.

For analyses on resolubility, we computed the average fraction of surface residues of a given type for all DODO (or chainsaw) domains associated with non-membrane proteins of a given resolubility quantile.

For analyses on structure retention, we mapped all the quantified peptides to DODO (or chainsaw) domains based on their residue boundaries. For tryptic peptides to be counted as part of a structural domain, >80% of the peptide’s residues had to fall within the domain boundary. Domains were divided by the fraction of their surface residues of a given type, and then the number of structurally perturbed and non-perturbed peptides associated with all domains in that category were counted. For the purpose of plotting, points are shown only for categories with ≥50 peptides coming from ≥5 proteins.

### Intermolecular interaction prediction

Surface-dependent intermolecular interactions were calculated using FINCHES, as described previously. FINCHES uses the physical chemistry encoded into molecular forcefields and repurposes that physical chemistry for bioinformatic analysis. We previously performed systematic clustering of all possible peptides to identify a minimal set of 36 distinct peptides that are chemically orthogonal to one another (see Fig. S11 in Ginell et al. (*35*)). For each individual DODO (or chainsaw) domain, we calculated the average mean-field attractive interaction between the surface of the domain and 36 distinct test peptides, enabling us to determine the “stickiness” of “inertness” of various domains.

### Evolutionary Domain-based Analyses

Proteins were divided up into domains based on the ECOD definitions <u>(Schaeffer et al. 2016)</u> and filtered using DomainMapper <u>(Manriquez-Sandoval and Fried 2022)</u>. For analyses of resolubility, we mapped the S/T resolubility value assigned to a given protein to all the constituent ECOD domains assigned to that protein and then analyzed the distribution of resolubility values for all domains associated with a given fold-type (or X-group). Data are provided in **Table S7**. For analyses on structure retention, we mapped all the quantified peptides to ECOD domains based on their residue boundaries. For tryptic peptides to be counted as part of a domain, >80% of the peptide’s residues had to fall within the domain boundary. Domains were divided by their fold-type, and then the number of structurally perturbed and non-perturbed peptides associated with all domains in that fold-type were counted. Significantly increased structural retention is defined as having fewer than half the expected number of perturbed peptides and a p-value < 0.01 by the chi-square test.

