## Supplemental Figures S1-S11 for "Protein surface chemistry encodes an adaptive tolerance to desiccation"

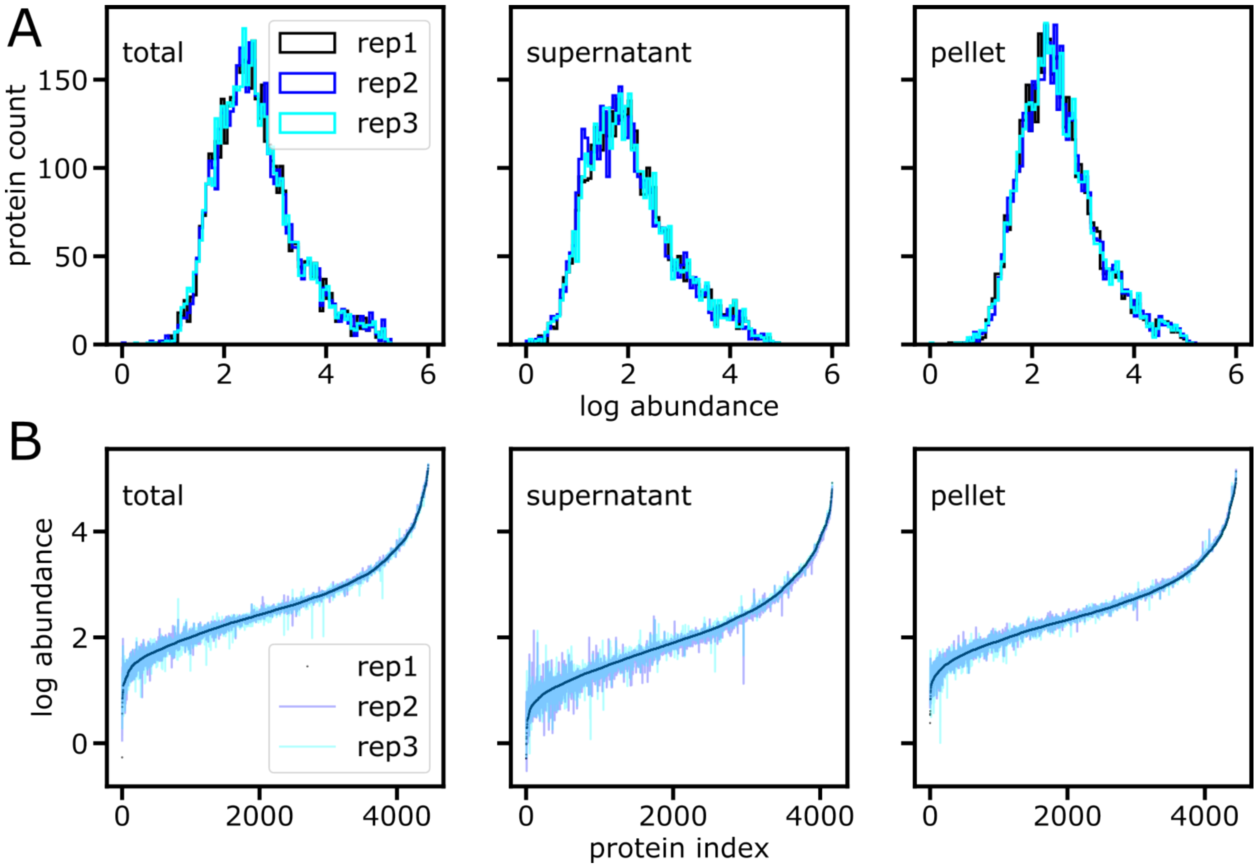


**Figure S1.** (**A**) Reproducibility of protein abundances (measured in ion counts, as calculated in Spectronaut) across triplicate samples for total, supernatant, and pellet. (**B**) Rank order reproducibility of protein abundances across triplicate samples for total, supernatant, and pellet.

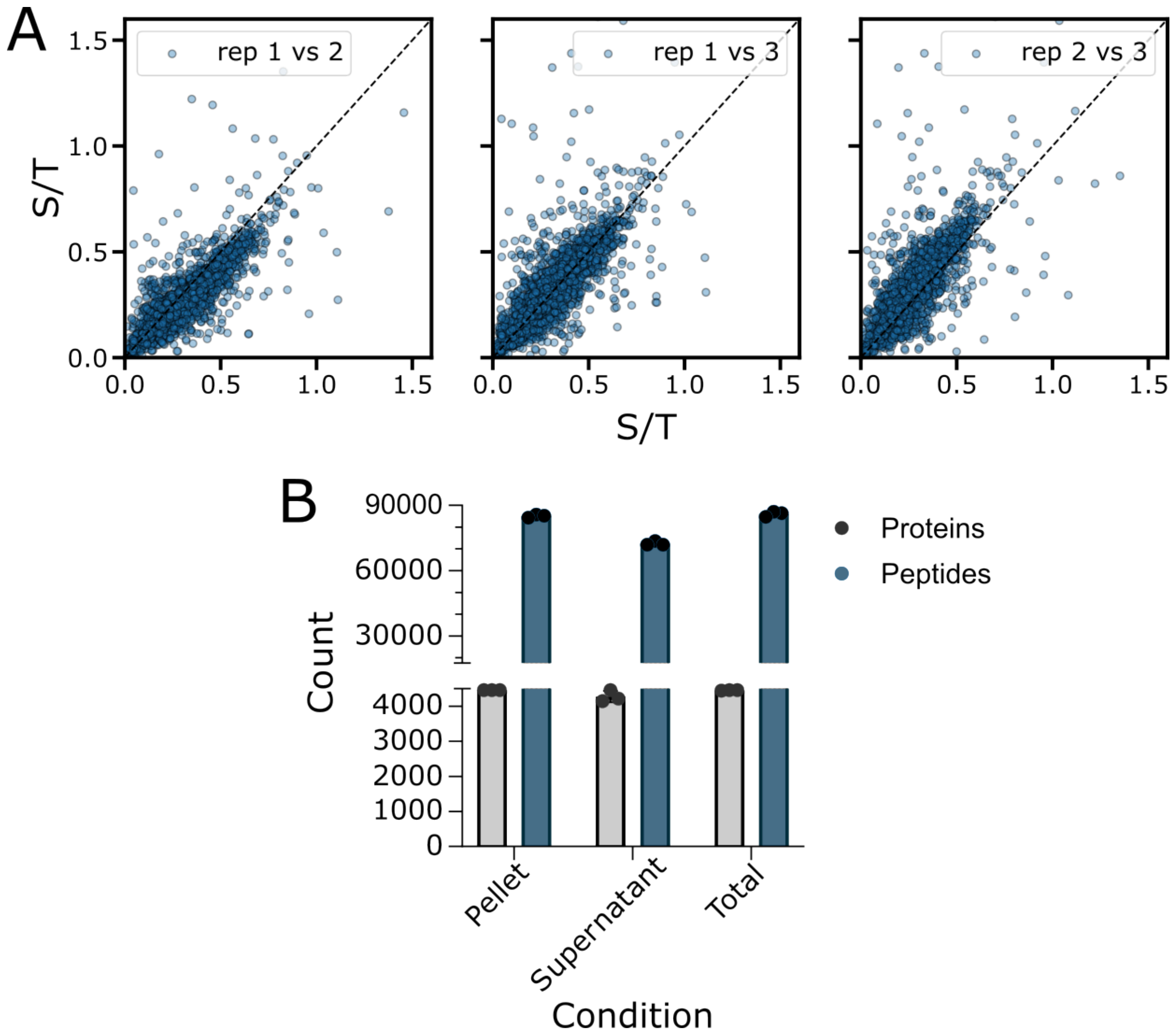


**Figure S2.** (**A**) Reproducibility of resolubility (as measured by S/T) across triplicate samples for proteome-wide resolubility experiment. (**B**) Number of identified proteins (grey) and peptides (blue) in total, supernatant, and pellet fractions from individual samples. The points represent triplicate biological repeats of the experiment.

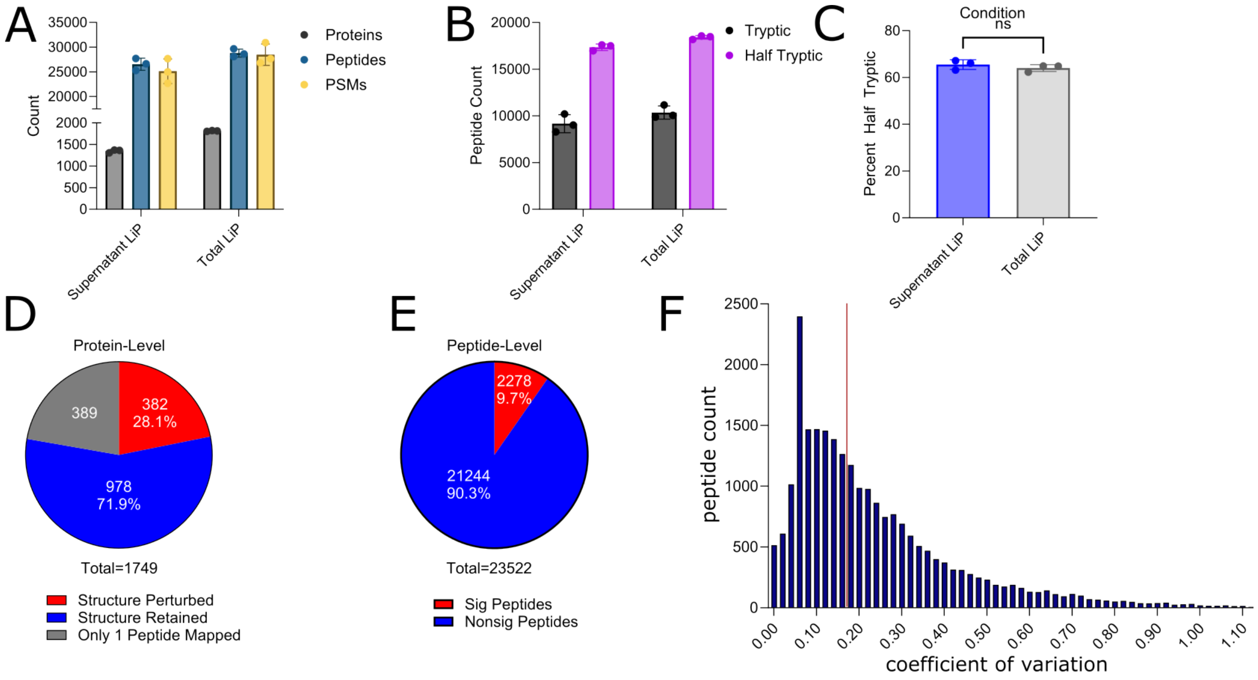


**Figure S3.** (**A**) Counts of proteins, peptides, and peptide-spectrum matches for individual samples from the limited proteolysis mass spectrometry (LiP-MS) experiment (where supernatant and total fractions were assessed). Points represent numbers from the three independent triplicates, and error bars are the standard deviation between the repeats. (**B**) Counts of identified tryptic and half-tryptic peptides from total and supernatant samples. Points represent numbers from the three independent triplicates, and error bars are the standard deviation between the repeats. (**C**) Percent of peptides that are half-tryptic from the overall number for total and supernatant samples. Points represent numbers from each of the three independent triplicates, and error bars are the standard deviation between the repeats. No significance was determined based on a student t-test. (**D**) Protein-level structure retention assignments from LiP-MS experiment. (**E**) Peptide-level structure retention assignments from LiP-MS experiment. (**F**) Distribution of peptide abundance coefficients of variation across biological replicates. The dashed red line denotes the median value.

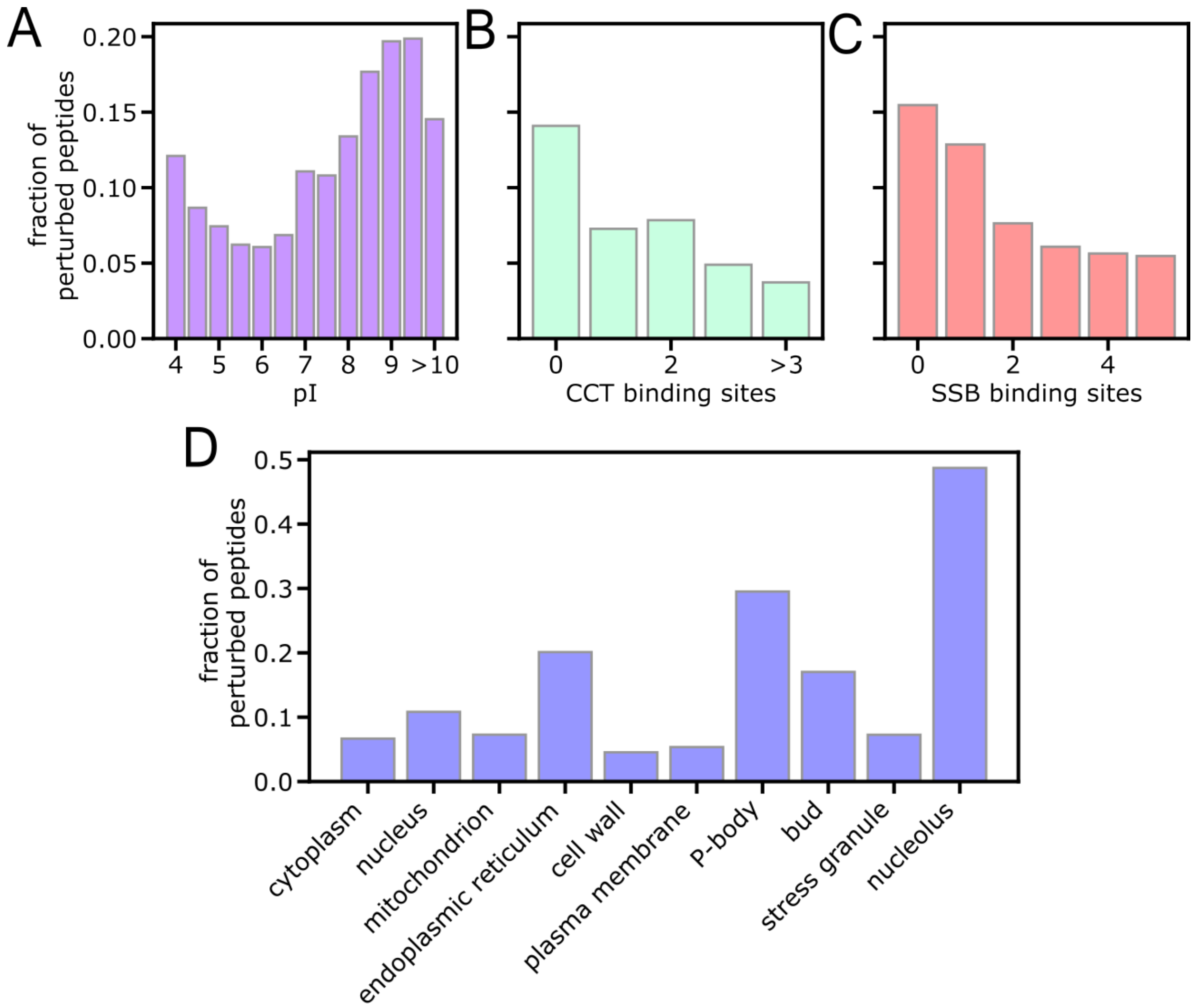


**Figure S4.** Fraction of peptides with structural alteration (change in proteolytic susceptibility) following desiccation-rehydration as a function of their parent proteins' (**A**) isoelectric point (pI), (**B**) number of CCT binding sites, (**C**) number of SSB binding sites, and (**D**) cellular location. Panels B and C use the number of CCT (i.e., chaperonin) and SSB (i.e., Hsp70) binding sites as measured by selective Ribo-Seq (Stein et al. 2019) In general we find that proteins which interact with chaperones more times during biogenesis exhibit greater structure retention (fewer perturbed peptides). A potential explanation for this finding is that chaperones may be more important for folding proteins with greater kinetic stability (or more rugged energy landscapes (To et al. 2022)), which could then in turn prevent structural disruption in the desiccated state.

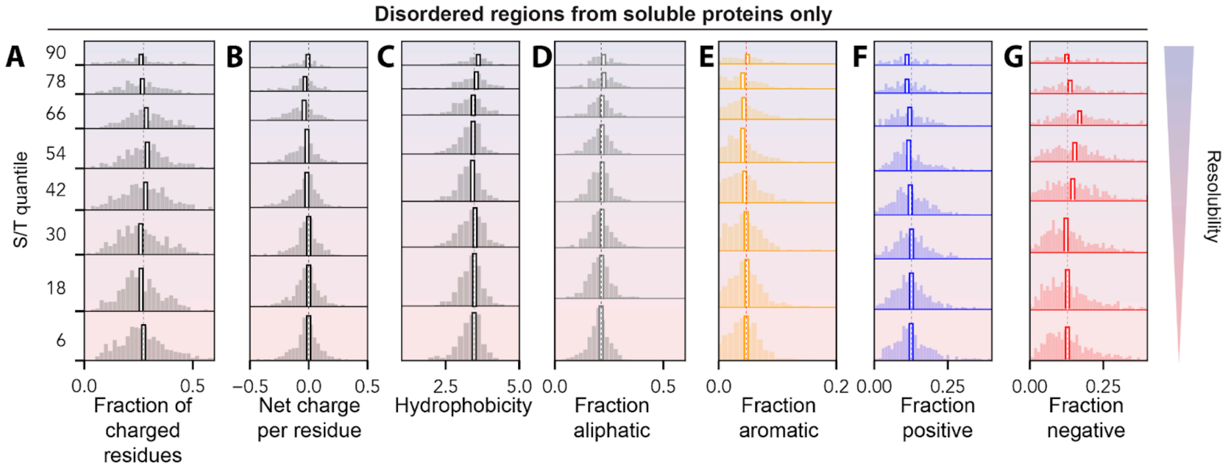


**Figure S5. Disordered protein correlations.** Disordered regions extracted from proteins (as defined by DODO) were assigned resolubility values and quantiles based on their parent proteins. For each disordered region, various residue-type fractions (and hydrophobicity) were calculated, and the distribution of those quantities is shown for all the disordered regions from all the proteins assigned to a given resolubility quantile. Distributions are shown for (**A**) fraction of charged residues (D + E + K + R); (**B**) net charge per residue (K + R - D - E); (**C**) hydrophobicity based on the average Kyte-Doolitle value; (**D**) fraction aliphatic (I + L + V + M); (**E**) fraction aromatic (W + Y + F); (**F**) fraction positive (K + R); and (**G**) fraction negative (D + E). Because resolubility quantities are assigned at the protein level, and more resoluble (higher S/T quantile) proteins have fewer disordered regions, the number of disordered regions associated with each histogram is not constant, but the number of proteins examined in each quantile is.

A

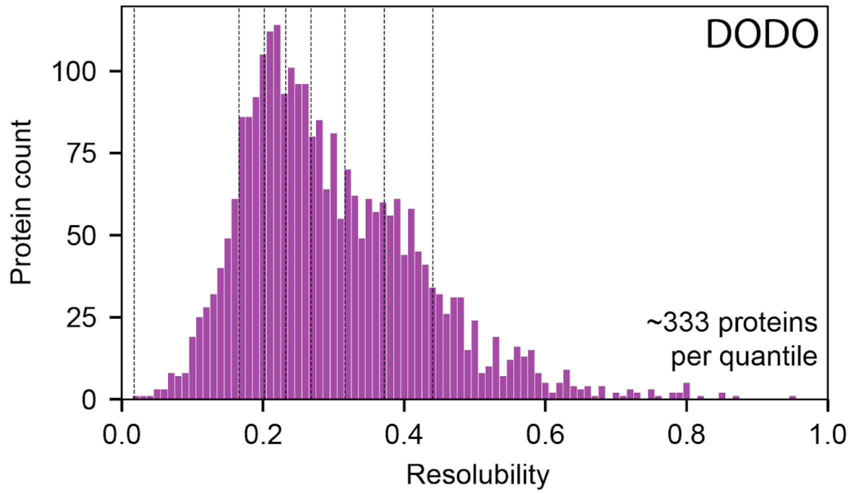


B

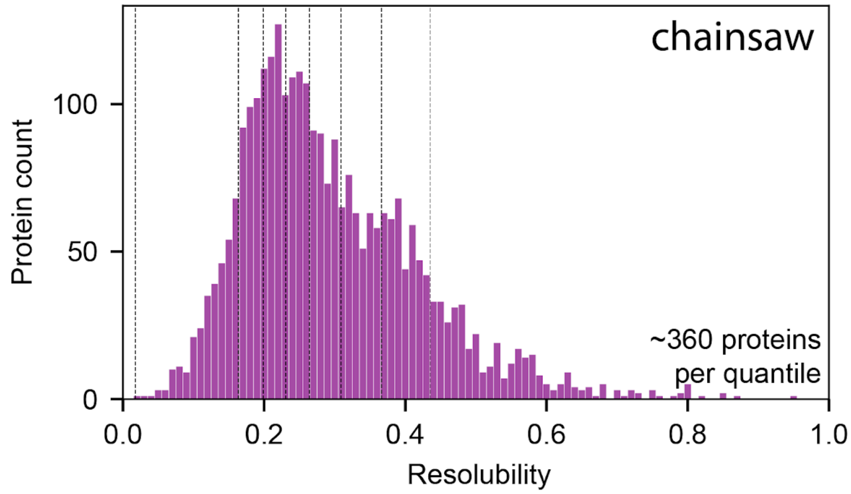


**Figure S6.** Distribution of the resolubilities of the structural domains of lengths greater than 90 amino acids (and less than 1000 amino acids) as defined by (A) DODO or (B) Chainsaw. Resolubility is defined as the mean S/T quantification. The measured resolubility for a given protein is assigned to all of its constituent structural domains. Eight quantiles are shown, demarcated by vertical dashed lines.

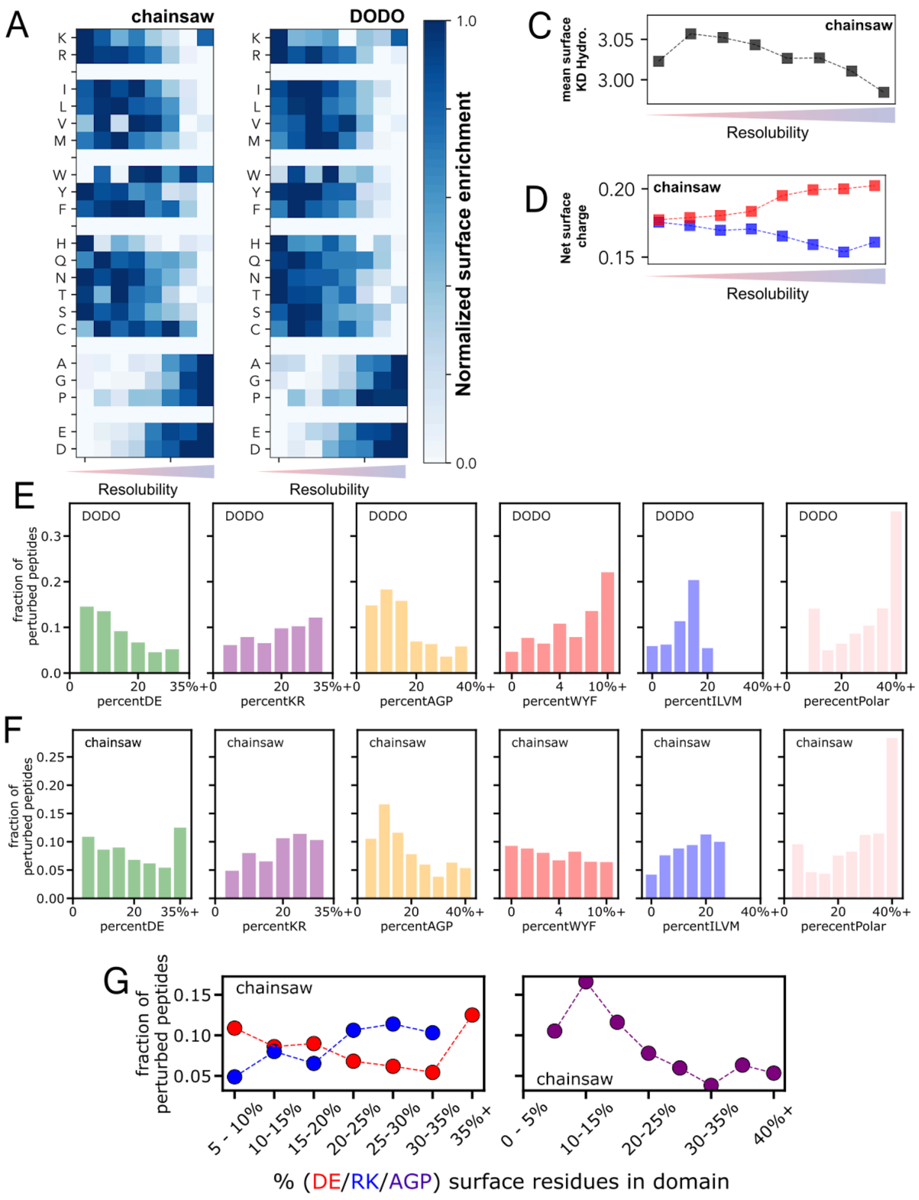


**Figure S7. Surface analysis comparison of chainsaw and DODO decomposition**. **(A-B)** Analysis of solvent-exposed amino acid fractions across different S/T quantiles using Chainsaw (A) and DODO-defined (B) domains. Each row is internally normalized such that each amino acid's surface fraction varies between 0 and 1, enabling comparison of amino acids with very different overall abundance. **(C-D)** Overall hydrophobicity (C) of surface-exposed residues and fraction of surface-exposed residues that are positive (blue, D) or negative (red, D) in different resolubility quantiles using chainsaw-defined domains. **(E-F)** Fraction of structurally perturbed peptides vs. the percentage of surface residues of the specified type according to DODO (E) and (F) Chainsaw-defined structural domains. **(G)** Fraction of structurally perturbed peptides from Chainsaw-defined domains with increasing percentages of positive (blue), negative (red), or small amino acids (purple) on the domain surfaces.

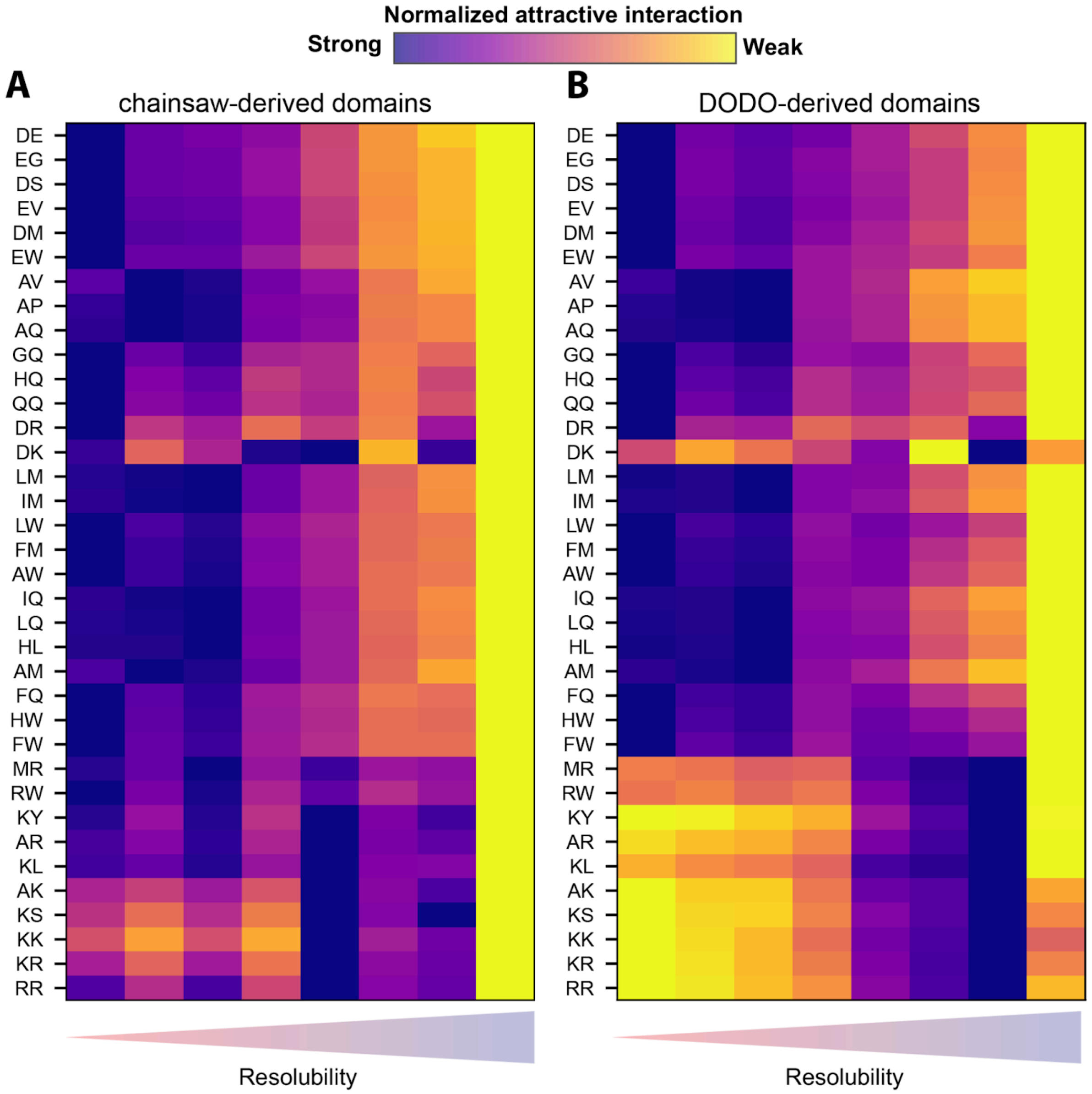


**Figure S8. A full comparison of all 36 chemically orthogonal dipeptides for domain surfaces defined using either Chainsaw or DODO.** The surfaces of structural domains compared to Chainsaw **(A)** and DODO **(B)** show comparable interaction energies with test peptides according to FINCHES. One exception is that DODO shows a more obvious clear skew in the positively-charged residues compared to chainsaw-derived domains, although in both cases, we see an enrichment of attractive interactions for positive residues in the more soluble proteins, consistent with the acquisition of negative charge. Attractive interaction scores are normalized within a given chemistry so all interactions can be plotted on the same axis.

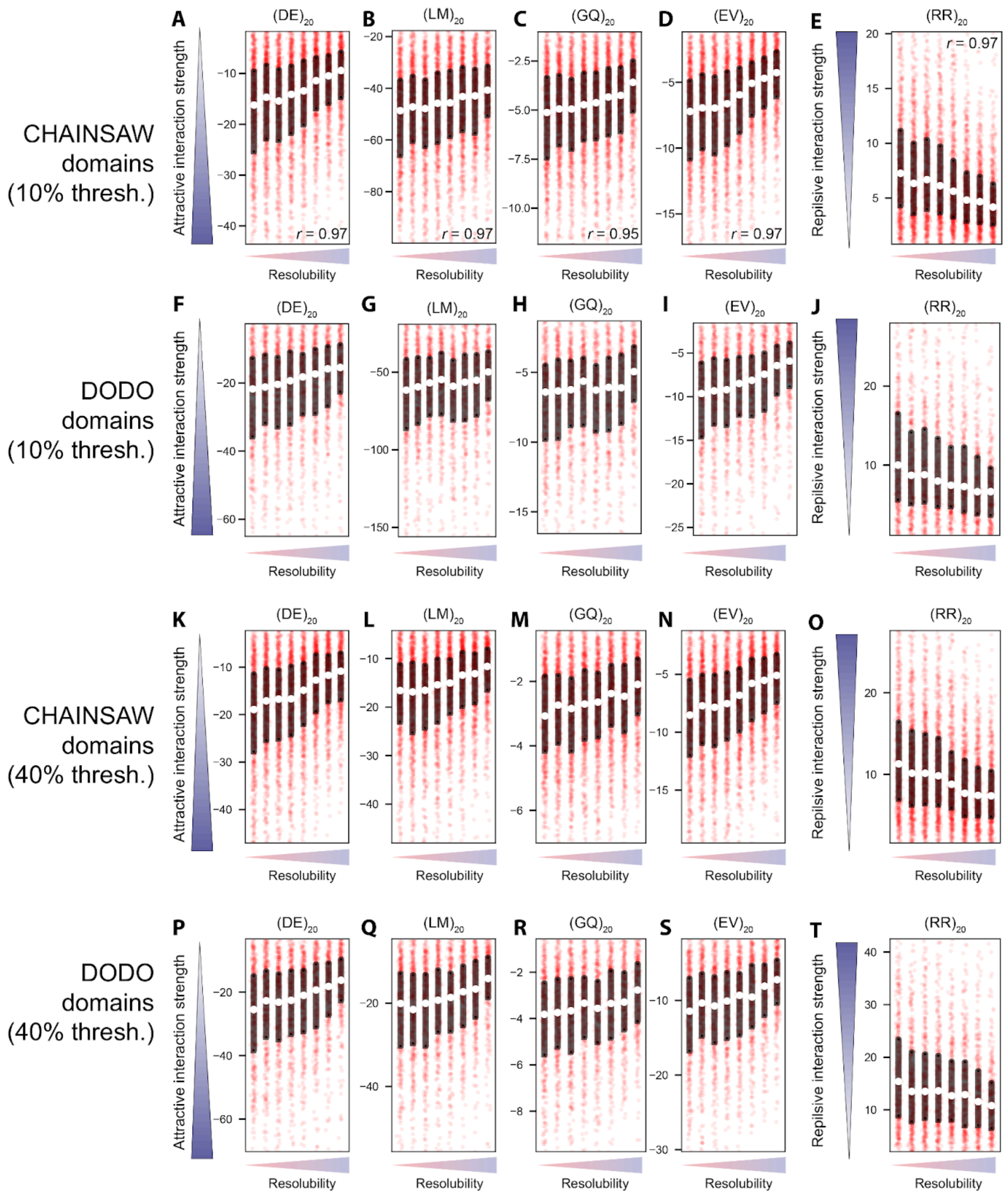


**Figure S9. Distribution of Dipeptide Interaction Strengths: Robustness Analysis.** Each panel corresponds to one of six representative test dipeptides whose interaction strength was predicted for each Chainsaw (panels A-E, K-O) and each DODO domain (panels F-J, P-T). Structural domains are divided by the resolubility quantile of the protein to which they belong, and the distribution of interaction scores for each quantile are shown. The use of DODO or Chainsaw with either a 10% threshold (panels A-E, F-J) or 40% threshold (panels K-O, P-T) on defining solvent-exposed residues has no impact on the overall trends observed.

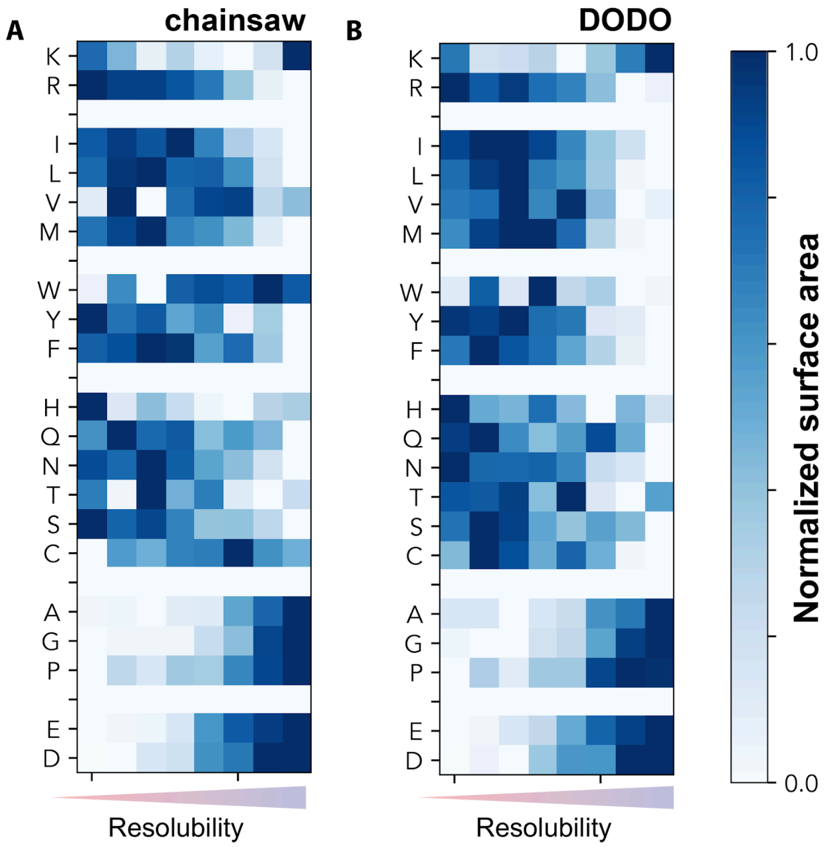


**Figure S10. Amino Acid Bias on the Surfaces of Resoluble Proteins: Robustness Analysis.** Analysis of amino acid enrichments on domain surfaces across different protein S/T quantiles according to (A) Chainsaw and (B) DODO defined decompositions. Each row is internally normalized such that each amino acid's surface fraction varies between 0 and 1, enabling comparison of amino acids with very different overall abundance. Unlike Fig. 4E, analysis of surface chemistry here uses normalized surface area instead of normalized amino acid fractions. Similar trends emerge as to those seen when normalized amino acid fractions are used.

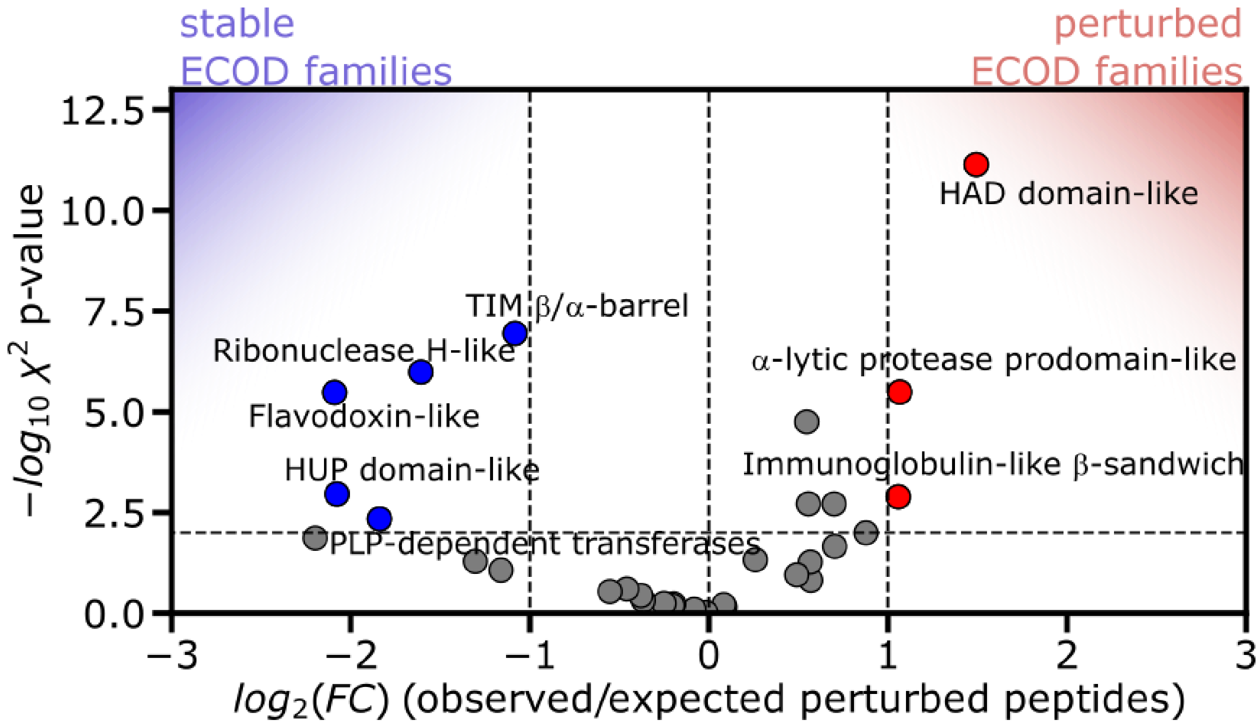


**Figure S11. Structure Retention of ECOD Domains.** Peptides were assigned to ECOD domains, and the total number of peptides (structurally perturbed or retained) of all domains of the same fold-type (X-group) were compiled. Volcano plot shows for each X-group the enrichment of structurally perturbed peptides (defined as the ratio of the observed to the expected number) and the p-value from the chi-square test against the null hypothesis that structurally perturbed peptides are not more frequent for that particular X-group. Dashed lines represent cutoffs for significance (>2-fold effect size, p-value < 0.01). Only folds with 100 or more peptides mapped to them are shown. Folds associated with metabolic function tend to exhibit structural retention: TIM barrels, ribonuclease H-like folds, flavodoxin-like folds, PLP-dependent transferase folds, and the HUP-domain (which includes class I aminoacyl-tRNA synthetases).
